# *Staphylococcus aureus* FtsZ and PBP4 bind to the conformationally dynamic N-terminal domain of GpsB

**DOI:** 10.1101/2022.10.25.513704

**Authors:** Michael D. Sacco, Lauren R. Hammond, Radwan E. Noor, Dipanwita Bhattacharya, Jesper J. Madsen, Xiujun Zhang, Shane G. Butler, M. Trent Kemp, Aiden C. Jaskolka-Brown, Sebastian J. Khan, Ioannis Gelis, Prahathees J. Eswara, Yu Chen

**Affiliations:** Department of Molecular Medicine, Morsani College of Medicine, University of South Florida, Tampa, FL, 33612, United States; Department of Cell Biology, Microbiology, and Molecular Biology, University of South Florida, Tampa, FL, 33620, United States; Department of Chemistry, University of South Florida, Tampa, FL, 33620, United States; Global and Planetary Health, College of Public Health, University of South Florida, Tampa, FL, 33612, United States

## Abstract

Bacterial cell division is a tightly regulated process that requires the formation of a dynamic multi-protein complex. In the Firmicutes phylum, GpsB is a membrane associated protein that coordinates peptidoglycan synthesis for cell growth and division. Although GpsB has been studied in several organisms, the structure, function, and interactome of *Staphylococcus aureus* GpsB is largely uncharacterized, despite being reported as uniquely essential for growth in this clinically relevant bacterium. To address this knowledge gap, we solved the crystal structure of the N-terminal domain of *S. aureus* GpsB. This structure reveals an atypical asymmetric dimer, and major conformational flexibility that can be mapped to a hinge region formed by a three-residue insertion exclusive to *Staphylococci*. When this three-residue insertion is excised, its thermal stability increases, and the mutant no longer produces a previously reported lethal phenotype when overexpressed in *Bacillus subtilis*. Furthermore, we provide the first biochemical, biophysical, and crystallographic evidence that the N-terminal domain of GpsB binds not only PBP4, but also FtsZ, through a conserved recognition motif located on their C-terminus, thus linking peptidoglycan synthesis with cell division. Taken together, the unique structure of *S. aureus* GpsB and its direct interaction with FtsZ/PBP4 provide deeper insight into the central role of GpsB in *S. aureus* cell division.

## Introduction

Bacterial cell division is a dynamic process involving more than a dozen proteins that form a multimeric complex at mid-cell. Collectively known as the divisome, this network of scaffolding proteins and enzymes stimulates peptidoglycan synthesis, constricts the existing membrane, and forms the septal cell wall^1^. While some proteins of the divisome differ amongst certain clades, most are conserved amongst all bacteria. Perhaps the most important and well-studied is FtsZ, a bacterial homolog of eukaryotic tubulin. FtsZ marks the division site by forming a “Z-ring” in association with early-stage divisomal proteins such as FtsA, ZapA, and EzrA^2,3^. Late-stage divisomal proteins such as DivIVA, FtsL, DivIB, and various penicillin-binding proteins (PBPs) subsequently assemble to carry out cell division and facilitate the separation and creation of identical daughter cells.

GpsB is a DivIVA-like protein that is highly conserved in Firmicutes^4,5^. While GpsB is dispensable, or conditionally essential in most Firmicutes^6–9^, it is reported to be essential for growth in *S. aureus*^10,11^. Notably, *S. aureus* GpsB (*Sa* GpsB) is unable to complement native GpsB in *Bacillus subtilis*, suggesting potential functional divergence. In fact, synthetic expression of *Sa* GpsB expression is lethal to *B. subtilis*^12^. The importance of *Sa* GpsB for cell division is underscored by its unique ability to regulate FtsZ polymerization in *S. aureus*^12^, in addition to interacting with other cell division proteins such as EzrA^13^, and the wall teichoic acids (WTA) biosynthesis/export proteins TarO and TarG^14,15^.

Several PBPs including *Listeria monocytogenes* (*Lm*) PBPA1, *Streptococcus pneumoniae* (*Sp*) PBP2a, and *B. subtilis* (*Bs*) PBP1 bind to GpsB through their N-terminal “mini-domains”; a sequence of ~ 5-30 residues on the cytosolic side of the cell membrane^6,16^. Recently, Cleverley et al. found PBP mini-domains containing an (S/T)-R-X-X-R-(R/K) motif directly interact with the N-terminal domain of GpsB by forming electrostatic interactions and hydrogen bonds with a shallow, acidic cavity located at the GpsB dimer interface (**Fig. S1**)^16^.

Whereas *Lm*, *Sp*, and *Bs* have at least six annotated PBPs^17–19^, there are only four in *S. aureus*: PBP1, PBP2, PBP3, and PBP4^11^ – five in methicillin-resistant *S. aureus* (MRSA) which expresses an additional β-lactam insensitive PBP, PBP2a^20^. *S. aureus* is further distinguished by its highly cross-linked peptidoglycan and can readily become resistant to β-lactams via PBP4 and the acquisition of PBP2a^21,22^. It is believed this β-lactam resistant phenotype relies on WTA assembly, which influences the function and localization of PBP4 and PBP2a^23^. While PBP2a is a historically recognized element of antibacterial resistance, PBP4 has recently been found to contribute to β-lactam insensitivity^24^. As the sole class C PBP in *S. aureus,* PBP4 bears the fold and architecture of a carboxypeptidase, but uniquely catalyzes both transpeptidase and carboxypeptidase reactions^25–28^.

In this report, we show that GpsB directly binds to FtsZ and PBP4 through a signature GpsB recognition sequence. Further analysis of the GpsB N-terminal domain reveals unique conformations and innate flexibility that is integral to the function of GpsB. Together, these findings provide insight to the unique role of GpsB in synchronizing FtsZ dynamics with cell wall synthesis during cell division in *S. aureus*.

## Results

### The crystal structure of *S. aureus* GpsB N-terminal domain reveals an atypical asymmetric dimer

The full-length *Sa* GpsB is a relatively small protein of 114 residues. Its N-terminal domain homodimerizes as a coiled-coil, while its smaller C-terminal domain homotrimerizes, forming a hexamer as the biological unit^4,6,29^. Using X-ray crystallography, we solved the structure of the N-terminal domain (res. 1-70) of *Sa* GpsB at 1.95 Å resolution in the P2_1_ space group (**Fig. 1A, Table S1**), with four monomers per asymmetric unit forming two dimers (dimer A and B). The overall structure of GpsB is similar to previously determined GpsB orthologues (**Fig. S1**)^6,16^ and retains the same fold of DivIVA^30^, though it shares much less sequence similarity (**Fig. 1E**). Dimerization of the N-terminal domain is facilitated by a pattern of nonpolar residues every three to four residues, promoting the formation of a hydrophobic core in the coiled-coil. This N-terminal domain is partitioned into three regions: a relatively short α-helix with approximately two turns (res. 10 - res. 16), a second longer α-helix (res. 28 - res. 70) that forms a coiled-coil, and an amphipathic ten-residue loop (res. 17 - res. 27) that links these two helices and intertwines with the adjacent protomer, “capping” GpsB (**Fig. 1B**). This loop region is proposed to interact with the inner leaflet of the cell membrane^31^.

**Figure 1.**
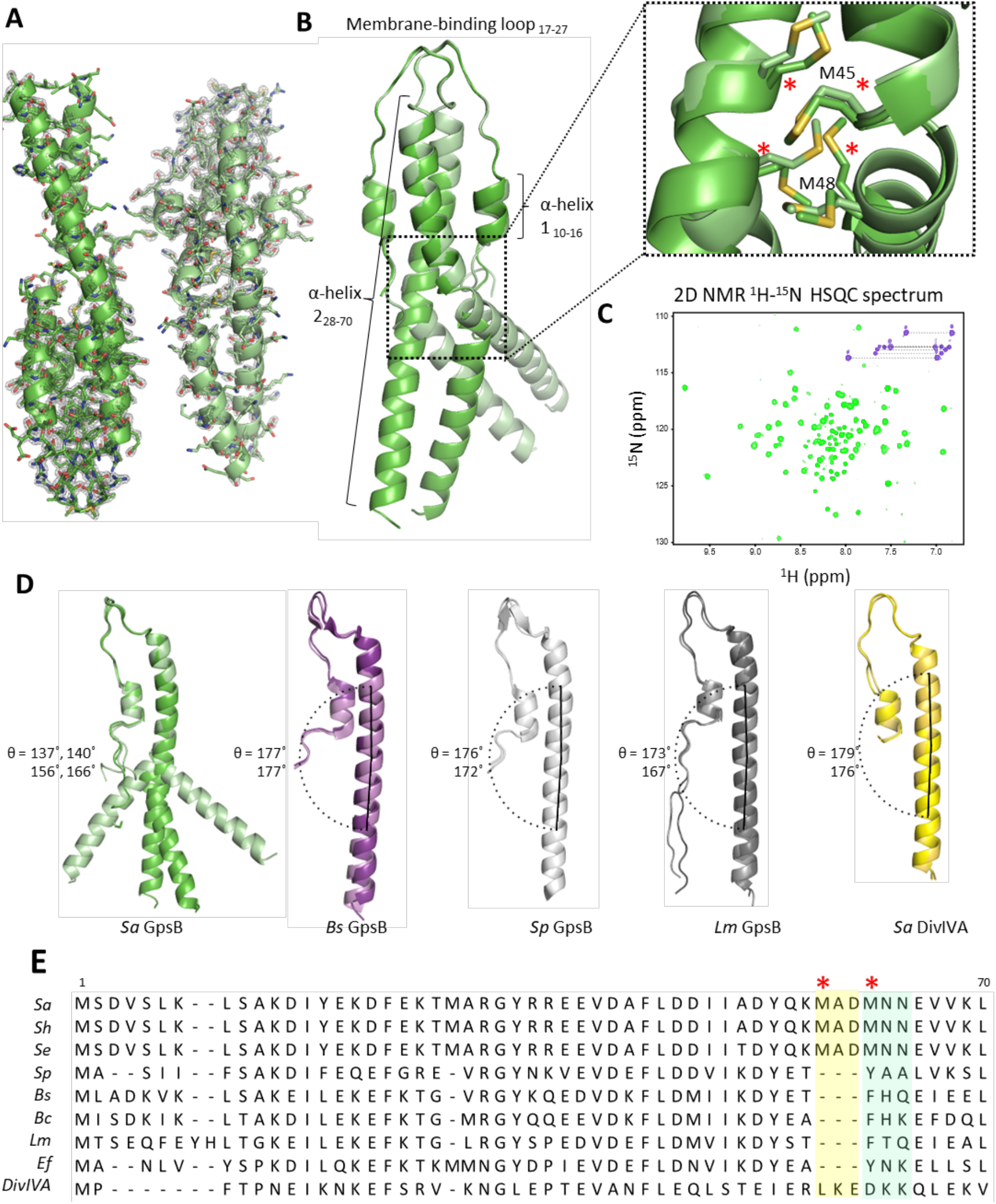
Crystal structure of the N-terminal domain of *S. aureus* GpsB. **(A)** Two dimers lie antiparallel in the asymmetric unit from the P2_1_ space group. The 2F_o_-F_c_ electron density map, shown in grey, is contoured at 1 σ with a resolution of 1.95 Å. **(B)** Superimposition of GpsB dimers reveals the dimers splay at a hinge region, formed by a cluster of four interlocked Met sidechains. *designates position in multisequence alignment shown in panel E. **(C)** The ^1^H-^15^N HSQC spectrum of the GpsB N-terminal domain, Sa *GpsB^WT^_1-70_*, shows a significantly higher number of signals compared to what is expected of a symmetric dimer based on the sequence of the domain, indicating the presence of conformational heterogeneity. Backbone and Asn/Gln sidechain amide signals are shown in green and purple respectively. **(D)** Comparison of different GpsB/DivIVA monomers from previously solved structures. Pitch angles were determined by placing a marker atom at the centroid of the top, bottom, and center turns of each helix, then measuring the angle. **(E)** Multisequence alignment of GpsB within select members of the Firmicutes phylum and with *S. aureus* DivIVA. *Staphylococci* GpsB contain a three-residue insertion that forms the hinge region, either MAD or MNN, depending on the sequence alignment parameters. The two Met residues (four per dimer) of the hinge region are designated with a red *. *Sa* - *S. aureus*, *Sh* -*S. haemolyticus*, *Se* - *S. epidermidis*, *Sp* - *S. pneumoniae*, *Bs* - *B. subtilis*, *Bc* - *B. cereus*, *Lm* - *L. monocytogenes, Ef* - *E. faecalis*, DivIVA - *S. aureus* DivIVA

The most unique structural feature of *Sa* GpsB is a hinge forming at the midpoint of its N-terminal domain that causes each protomer to bend, adopting pitch angles (θ) of 137°-140° (dimer A), and 156° - 166° (dimer B). In contrast, GpsB protomers from *Lm, Bs*, *Sp*, and *Sa* DivIVA are almost linear (θ = 167° - 179°) and practically indistinguishable when superimposed (**Fig. 1D**). We find that there is a 3-amino acid insertion in the *S. aureus* GpsB sequence where the helicity is disrupted and that a cluster of four Met residues (Met45, Met48) are interlocked at the dimer interface at this position (**Fig. 1B**; dotted rectangle). These Met residues are only found in *Staphylococci* (**Fig. 1E**) and take the place of an aromatic Tyr or Phe present in other orthologs, which normally stabilize the core via π stacking interactions. While the 3-aa insertion likely disrupts the continuity of the coiled-coil, methionine is one of the most flexible aliphatic amino acids and may further contribute to the conformational flexibility.

Experiments using solution state NMR spectroscopy further support the notion of intrinsic conformational heterogeneity, suggesting it is a *bona fide* structural feature rather than a crystallization artifact. The *Sa* GpsB^WT^_1-70_ construct contains a two-residue N-terminal extension (GH) after removal of the purification tag, and thus a total of 72 signals are expected. The 2D ^1^H-^15^N HSQC spectrum shows good signal dispersion **(Fig. 1C)**, which is characteristic of a folded domain. However, a total of 100 well-resolved backbone amide signals are observed, and seven pairs of signals are detected in the Asn/Gln sidechain region instead of four pairs **(Fig. 1C, Fig. S2A**). The appearance of the additional signals is not due to proline *cis/trans* isomerization since there are no prolines in the sequence, nor is it due to self-aggregation of the dimers to form a dimer-of-dimers or higher order oligomers, because raising the concentration from 160 μM to 700 μM has no effect on the number of signals in the spectrum (**Fig. S2B**). In addition, the ^1^H-^15^N HSQC of *Sa* GpsB^WT^_1–70_ exhibits differential linewidths, with a set of 13 narrow signals at the center of the spectrum where disordered segments appear, and a larger set of 87 signals dispersed throughout the spectrum. The number of the narrow signals corresponds well with the number of residues found at the flexible N-terminus, suggesting that the appearance of extra signals is due to sampling of multiple conformations of the coiled-coil. Furthermore, three of the four Asn/Gln sidechain pairs showing chemical shift degeneracy are adjacent to the hinge **(Fig. S2A)** providing strong evidence that the observed conformational heterogeneity occurs at this region.

MD simulations using Gromacs v. 5.0.4 and a CHARMM36m force field also support the idea that conformational flexibility exists in solution. After approximately 100 ns, dimer 1 adopts an ~180° pitch angle that occurs concomitantly to major fluctuations in dimer 2 in a separate simulation **(Fig. S3A)**, although the preference for continuous helical structures (corresponding to ~180° pitch angles) may sometimes be influenced by specific force field parameters.

### A three-residue insertion unique to *Staphylococci* GpsB disrupts the coiled-coil pattern and destabilizes the structure of *Sa* GpsB

A unique element of *Sa* GpsB that may contribute to its distinct conformational flexibility are three extra residues located at the hinge region– MAD or MNN (depending on the alignment parameters; **Fig. 1E**), roughly corresponding to an extra turn in the helix. Deleting either MAD or MNN produces a homology model where residues 45-70, normally displaced by one turn, are now aligned with their orthologous residue pair (**Fig. S3B**). To investigate the relative degree of stability imparted by these residues, they were genetically excised, recombinantly expressed, and their T_m_ (melting temperature) was determined using circular dichroism (CD) spectroscopy (**Fig. 2A**). This experiment shows the ΔMAD and ΔMNN GpsB have superior thermal stability to WT *Sa* GpsB for both the N-terminal domain (1-70) and full-length constructs. Perhaps the most notable finding from this experiment was that, while the T_m_ of *Sa* GpsB^ΔMAD^_1-70_ (38.88 ± 0.18 °C) was only modestly higher than *Sa* GpsB^WT^_1-70_ (34.63 ± 0.33 °C), the T_m_ of *Sa* GpsB^ΔMNN^_1-70_ (53.65 ± 0.55 °C) was highly stable, approximately 1.5-fold higher than that of *Sa* GpsB^WT^ _1-70_ and comparable to the thermal stability of the full-length *Sa* GpsB^WT^ _FL_. Though the full-length *Sa* GpsB^ΔMNN^ _FL_ was not analyzed, we expect that its T_m_ would be higher than both *Sa* GpsB^WT^ _FL_ and *Sa* GpsB^ΔMAD^ _FL_ based on the experiments assessing the 1-70 constructs.

**Figure 2.**
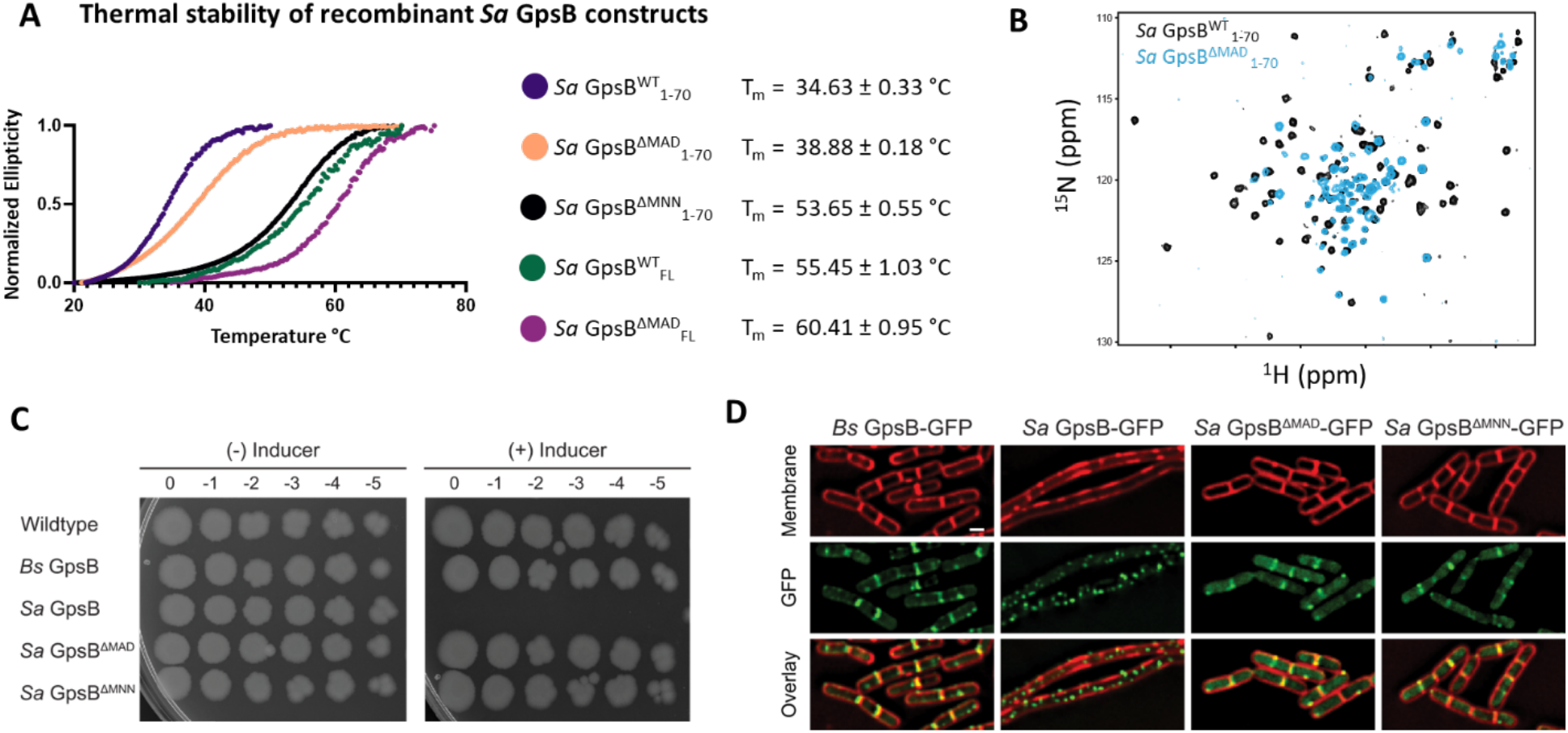
Deletion of a three-residue insertion in *Sa* GpsB increases thermal stability in solution and abolishes toxicity in *B. subtilis*. **(A)** CD melt profiles of recombinantly expressed *Sa* GpsB constructs reveal ΔMAD and ΔMNN mutants have increased T_m_, compared to WT *Sa* GpsB. **(B)** Overlays of the ^1^H-^15^N HSQC of spectrum of *Sa* GpsB^WT^ _1-70_ and *Sa* GpsB^ΔMAD^ _1-70_ suggests there are significant differences in the conformational properties of these two proteins. **(C)** Serial dilutions of *B. subtilis* strains harboring inducible *Bs* GpsB (GG18), *Sa* GpsB (GG7), *Sa* GpsB ΔMAD (LH119), and *Sa* GpsB ΔMNN (LH115), plated on LB plates without (left) and with (right) 1 mM IPTG demonstrate WT *Sa* GpsB is lethal, but ΔMAD *Sa* GpsB and ΔMNN *Sa* GpsB are not. **(D)** Fluorescence micrographs showing the protein localization of *Bs* GpsB-GFP (GG19), *Sa* GpsB-GFP (GG8), *Sa* GpsB-GFP ΔMAD (LH126), and *Sa* GpsB-GFP ΔMNN (LH116). Cell membrane was visualized using SynaptoRed membrane dye (1 μg/ml). Scale bar, 1 μm. In contrast to WT *Sa* GpsB, strains of *B. subtilis* that overexpress ΔMAD and ΔMNN *Sa* GpsB have similar cellular morphology to WT *B. subtilis* and these proteins localize to the division septum.

The conformational properties of the *Sa* GpsB^ΔMAD^ _1-70_ mutant were also different when analyzed by ^1^H-^15^N HSQC (**Fig. 2B**). Most of the dispersed signals from the coiled coil region experience severe broadening, in many cases beyond detection, as well as changes in chemical shift, while all narrow signals at the center of the spectrum show no change in linewidth or positions. Signal broadening is caused by changes in the rate of interconversion between the available conformations from the intermediate-slow for WT (seconds) to the intermediate-fast exchange regime for ΔMAD (milliseconds), but without altering the overall number of states, as seven pairs of Asn/Gln sidechain pairs of signals are observed. Conformational rigidity is a well-established correlate of thermal stability^32^, and may contribute to the enhanced T_m_ of *Sa* GpsB^ΔMNN^ and *Sa* GpsB^ΔMAD^. The aforementioned homology model **(Fig. S3B)** corresponding to *Sa* GpsB^ΔMNN^ and *Sa* GpsB^ΔMAD^ reveals several residues potentially form stronger interactions than the WT. These include an intrahelical electrostatic interaction that replaces a potential repulsion between Asp47 and Glu51 with Asn47/Lys51 in *Sa* GpsB^ΔMAD^ and Asp47/Lys51 in *Sa* GpsB^ΔMNN^ (**Fig. S3C**). The favorable Asp47/Lys51 electrostatic interactions in GpsB^ΔMNN^ may further contribute to its higher thermostability.

### The MAD/MNN insertion is critical for GpsB function

Previously, we reported that overproduction of *Sa* GpsB in *B. subtilis* causes cell division arrest which eventually leads to filamentation and cell lysis^12^. We used this system to probe the significance of the flexibility provided by MAD/MNN residues for the function of GpsB. First, we conducted a growth assay on solid medium by spotting serial dilutions of cells of *B. subtilis* wildtype (WT) and cells harboring inducible copy of *Bs* GpsB, *Sa* GpsB, *Sa* GpsB^ΔMAD^, or *Sa* GpsB^ΔMNN^. As previously established, overproduction of *Bs* GpsB is not lethal, but *Sa* GpsB is (**Fig. 2C**)^12,15^. In contrast, overproduction of either ΔMAD or ΔMNN *Sa* GpsB is not lethal and cells grow as well as the negative controls (*B. subtilis* WT and inducible *Bs* GpsB strain). These results suggest the hinge region of *Sa* GpsB demarked by Met residues (**Fig. 1B**) is essential for the normal function of *Sa* GpsB as assessed by lethal phenotype in *B. subtilis*. To further investigate the impact of these mutations, we examined GFP tagged ΔMAD and ΔMNN *Sa* GpsB in *B. subtilis* using high resolution fluorescence microcopy **(Fig. 2D)**. As reported previously, and as shown in **Fig. 2D**, *Bs* GpsB-GFP localizes to the division site^8,9,15^ and does not lead to filamentation upon overproduction (2.09 μm ± 0.49 μm; n=50), but *Sa* GpsB-GFP localizes, forms foci throughout the entire cell, and causes severe filamentation (22.83 μm ± 16.51 μm; n=21)^12^. However, ΔMAD and ΔMNN *Sa* GpsB-GFP do not cause filamentation (2.13 μm ± 0.44 μm and 2.10 μm ± 0.43 μm respectively; n=50) and are clearly localized at the division site. It has been noted previously that *Sa* GpsB-GFP also localizes at division sites at initial stages prior to causing filamentation at a lower inducer concentration^12^. Taken together, this data suggests the ΔMAD and ΔMNN mutants presumably interact with the *B. subtilis* cell division machinery to allow for division site localization, but fail to elicit lethal filamentation and toxicity^12^.

We also investigated the phenotypes of ΔMAD and ΔMNN *Sa* GpsB in *S. aureus*. First, we conducted a growth assay by spotting serial dilutions of *S. aureus* strains harboring an empty vector (EV) or an additional plasmid-based inducible copy of *Sa gpsB*, *Sa gpsB^ΔMAD^*, or *Sa gpsB^ΔMNN^*. Overproduction of *Sa* GpsB leads to a 100 to 1000-fold growth inhibition compared to EV control (**Fig. S4A**). Interestingly, while cells overproducing *Sa* GpsB^ΔMNN^ grew similar to the EV control, *Sa gpsB^ΔMAD^* overexpression resulted in growth inhibition similar to *Sa* GpsB. We hypothesized that the difference in phenotype is due to differential affinity of native *Sa* GpsB (produced from chromosomal locus) to ΔMAD or ΔMNN mutants (produced from a plasmid-based system), leading to less or more propensity for heterocomplex formation. To test this, we conducted bacterial two-hybrid analysis as reported previously^15,33^. As shown in (**Fig. S4B**), the affinity of *Sa* GpsB^ΔMNN^ to itself appears to be greater compared to *Sa* GpsB^ΔMNN^ and *Sa* GpsB. *Sa* GpsB^ΔMAD^ appears to have similar affinity to itself and *Sa* GpsB, however it is lower compared to the self-interaction of *Sa* GpsB. Thus, the differential affinity between mutants and *Sa* GpsB is the likely source of different phenotypes. Second, we wanted to analyze the cell morphology of *S. aureus* strains overproducing ΔMAD/ΔMNN mutants, as we have previously shown overproduction of *Sa* GpsB leads to cell size enlargement due to cell division inhibition^12^. Using fluorescence microscopy (**Figs. S4C** **and** **S4D**), we re-confirmed the increase in cell diameter in cells overproducing *Sa* GpsB (1.00 μm ± 0.19 μm) when compared to the EV control (0.90 μm ± 0.13 μm). In agreement with the lethal plate phenotype (**Fig. S4A**), cells overproducing *Sa* GpsB^ΔMAD^ also displayed a statistically significant increase in cell diameter (0.95 μm ± 0.16 μm), while *Sa* GpsB^ΔMNN^ did not (0.91 μm ± 0.15 μm) and resembled EV control. We also ensured the stable production of *Sa* GpsB, ΔMAD, and ΔMNN via western blotting (**Fig. S4E**). Lastly, by GFP tagging, we observed that both mutants localize to the division site similar to *Sa* GpsB-GFP (**Fig. S4F**)^12,15^. In summary, ΔMNN is less functional compared to ΔMAD in terms of causing growth inhibition on plate and cell enlargement phenotype, however both ΔMAD and ΔMNN mutants are able to localize to division sites.

### The C-terminus of *Sa* FtsZ binds to the N-terminal domain of *Sa* GpsB through a conserved (S/T/N)-R-X-X-R-(R/K) motif

One of the few proteins known to interact with *Sa* GpsB is the tubulin-like GTPase, FtsZ - a central cell division protein that marks the division site in nearly all bacteria. We previously demonstrated that GpsB directly interacts with *Sa* FtsZ to stimulate its GTPase activity and modulate its polymerization characteristics^12^. However, the molecular basis for this interaction was not known. Remarkably, we discovered that the last 12 residues of *Sa* FtsZ (N-R-E-E-R-R—S-R-R-T-R-R), also known as the C-terminal variable (CTV) region^34^, are a repeated match of the consensus GpsB-binding motif (S/T-R-X-X-R-(R/K)) found in the N-termini of *Bs* PBP1, *Lm* PBPA1, and *Sp* PBP2a. In this instance, the first motif bears an Asn instead of a Ser/Thr. Given the similar physicochemical properties of Asn as a small polar amino acid, it can likely replace Ser or Thr without any functional significance. To our knowledge, the interaction between GpsB and FtsZ is unique to *S. aureus^5^*, which is consistent with the absence of this motif in FtsZ orthologs from other organisms (**Fig. 3A**).

**Figure 3.**
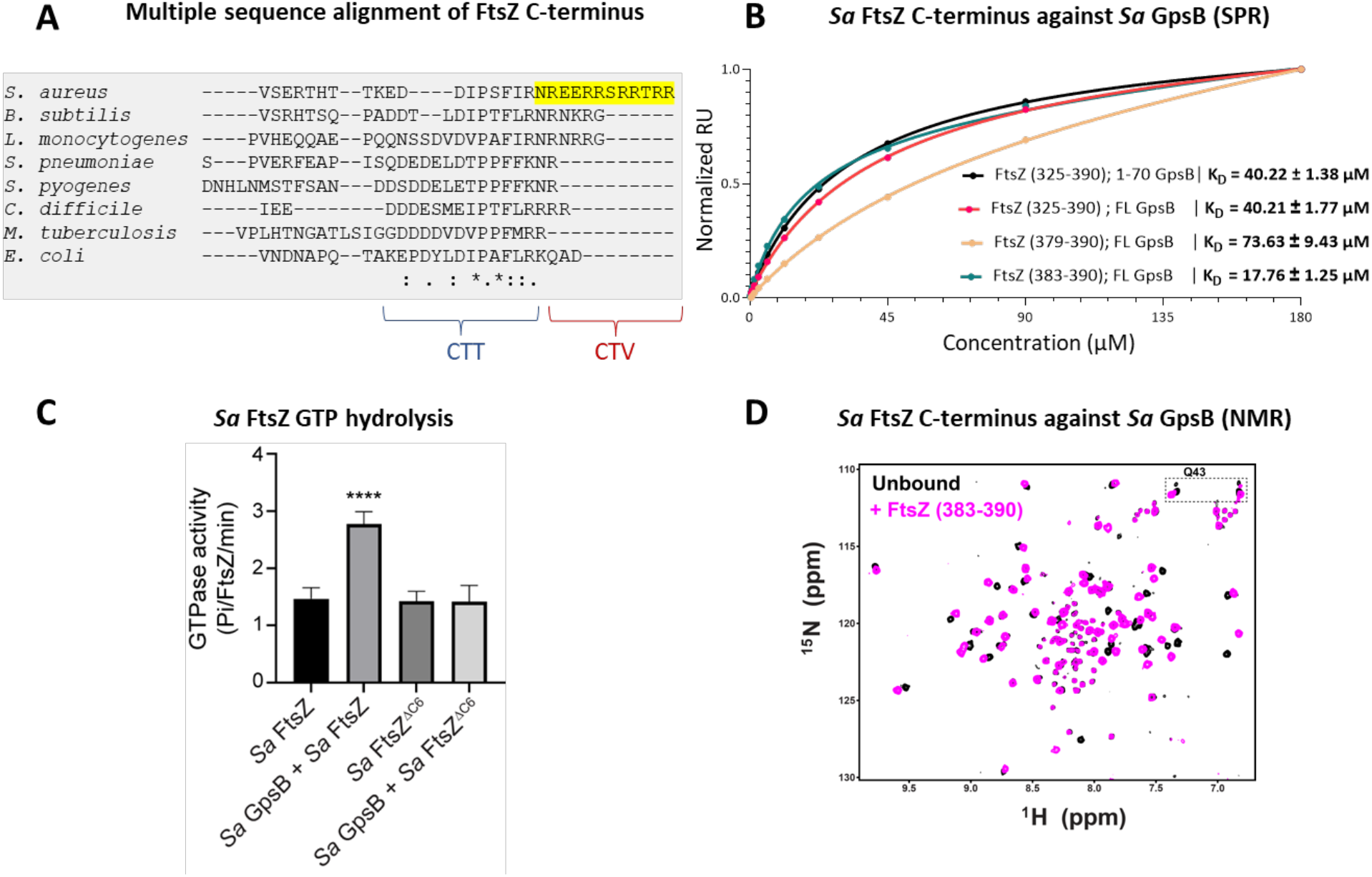
*Sa* FtsZ contains a repeated GpsB-recognition motif at its C-terminus. **(A)** A multiple sequence alignment of the FtsZ C-terminus from different representative bacteria reveals that there is a repeated GpsB-recognition motif in *Sa* FtsZ (highlighted region) and that it is unique to this bacterium. **(B)** SPR titration of peptides corresponding to several segments of *Sa* FtsZ against *Sa* GpsB (residue 1-70 or full-length). A titration of Sa FtsZ (res. 325-390) against 1-70 GpsB (black), corresponding to the N-terminal domain, shows binding can be isolated to this region. **(C)** When incubated with *Sa* GpsB, a *Sa* FtsZ mutant with a C-terminal truncation (SRRTRR, 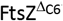) has significantly lower GTP hydrolysis compared to its full-length counterpart. GTP hydrolysis was measured by monitoring inorganic phosphate (P_i_) released (μmoles/min) by either FtsZ or FtsZ^ΔC6^ (30 μM) in the absence and presence of GpsB (10 μM). The plot is the average of n=6 independent data sets. P value for **** is < 0.0001. **(D)** Overlays of the ^1^H-^15^N HSQC of spectrum of *Sa* GpsB (1-70) in the absence (black) and in the presence of FtsZ (383-390; pink). The boxed region highlights the only sidechain pair of signals that becomes affected by the addition of Ftsz. Based on a model derived from the structure of Bs GpsB in complex with a PBP-derived peptide (Fig. S2), it is tentatively assigned to Q43 of Sa GpsB.

To initially test whether the predicted FtsZ GpsB-binding motif is actually involved in binding to *Sa* GpsB, we purified and titrated the terminal 66 residues of *Sa* FtsZ (325-390) against full-length *Sa* GpsB using surface plasmon resonance (SPR), revealing a dose-dependent interaction (K_D_ = 40.21 ± 1.77 μM; **Fig. 3B**). A simultaneous titration against only the *Sa* GpsB N-terminal domain (1-70) demonstrates that the C-terminus of *Sa* FtsZ binds exclusively to this domain (K_D_ = 40.22 ± 1.38 μM). This is further supported by assays with *Sa* FtsZ CTV (379-390; K_D_ = 73.63 ± 9.43 μM) and the final eight residues of *Sa* FtsZ (383-390; K_D_ 17.76 ± 1.25 μM). The described biophysical affinity aligns with previous cellular studies^16^, and is consistent with our hypothesis that the impetus for binding is the (S/T/N)-R-X-X-R-(R/K) motif which associates with the PBP-binding pocket (**Fig. S1**). Unlike the previously identified GpsB binding motifs located at the N-termini of PBPs, this is the first time such a motif has been found at the C-terminus of a GpsB-binding protein. The approximate four-fold difference in affinity between the CTV and the final eight residues could be a result of the inclusion of two Glu residues in the CTV (E381, E382), which may experience repulsion from the negatively charged PBP-binding pocket of GpsB. Notably, the GpsB recognition motif of *Sa* FtsZ is adjacent to the C-terminus carboxylate, which imparts an additional negative charge.

To further characterize the interaction between *Sa* FtsZ and *Sa* GpsB, we conducted a GTPase assay with *Sa* GpsB and *Sa* FtsZ or *Sa* FtsZ with the C-terminal six residues truncated (*Sa* FtsZ^ΔC6^). Briefly, in our previous report^12^, we found that *Sa* GpsB enhances the GTPase activity of *Sa* FtsZ. Therefore, we hypothesized that truncation of the terminal six residues of *Sa* FtsZ would eliminate the GpsB-mediated enhancement of GTPase activity. As shown in (**Fig. 3C)**, and as reported previously, addition of *Sa* GpsB enhanced the GTPase activity of *Sa* FtsZ. However, this effect was not seen in *Sa* FtsZ^ΔC6^ suggesting that the last six C-terminal residues of *Sa* FtsZ is likely where the interaction with *Sa* GpsB occurs.

The interaction of the *Sa* GpsB N-terminal domain with the *Sa* FtsZ derived peptide (383-390) was also monitored using NMR (**Fig. 3D**). Addition of the octapeptide to ^15^N-labeled N-terminal domain results in a significant chemical shift perturbation to a small number of dispersed signals in the ^1^H-^15^N HSQC spectrum, suggesting that the interaction is highly localized and occurs through the coiled-coil region without disturbing the conformational heterogeneity of the dimer. The available structures of *Bs* GpsB and *Sp* GpsB in complex with PBP-derived peptides (**Fig. S1 A, C**) suggest the complex is stabilized through interactions with helices 1 and 2, as well as part of the h1-h2 capping loop^16^. The N-terminal Arg of *Sa* FtsZ octapeptide utilized in our experiments is expected to be placed in the PBP-binding pocket near Q43, based on the complex structures of GpsB homologs. Indeed, only one of the Asn/Gln pairs of signals in the ^1^H-^15^N HSQC is shifted upon addition of the peptide, while in agreement with our model all other sidechain signals lie far from the binding site (**Fig. S2A**), suggesting that *Sa* GpsB recognizes partner proteins in a conserved manner.

### The cytosolic, C-terminal mini-domain of *Sa* PBP4 binds to *Sa* GpsB through its (N)-R-X-X-R-(R) recognition motif

*S. aureus* has an unusually low number of PBPs in its genome^11^. Although PBP1, PBP2, PBP2a, and PBP3 all have a cytoplasmic N-terminal “mini-domain”, none bear a GpsB recognition motif **(Fig. 4A)**. Using SPR, we confirmed their cytoplasmic mini-domains have no affinity for *Sa* GpsB, which is in agreement with the findings from previous bacterial two-hybrid assays^13^. Remarkably, PBP4, the only class C PBP encoded in the *S. aureus* genome, which purportedly functions as both a carboxypeptidase and transpeptidase PBP^35^, has a short, cytosolic C-terminal mini-domain with the sequence of N-R-L-F-R-K-R-K, satisfying the consensus GpsB-binding motif (S/T/N)-R-X-X-R-(R/K) found in *S. aureus* FtsZ and in orthologous PBPs found to bind to GpsB. Next, using SPR, we demonstrate this *Sa* PBP4 C-terminal octapeptide binds to *Sa* GpsB with a K_D_ of 48.61 ± 1.86 μM **(Fig. 4A)**. Supporting this finding is a potential interaction between *Sa* PBP4 and *Sa* GpsB previously noted in a bacterial two-hybrid study^14^.

**Figure 4.**
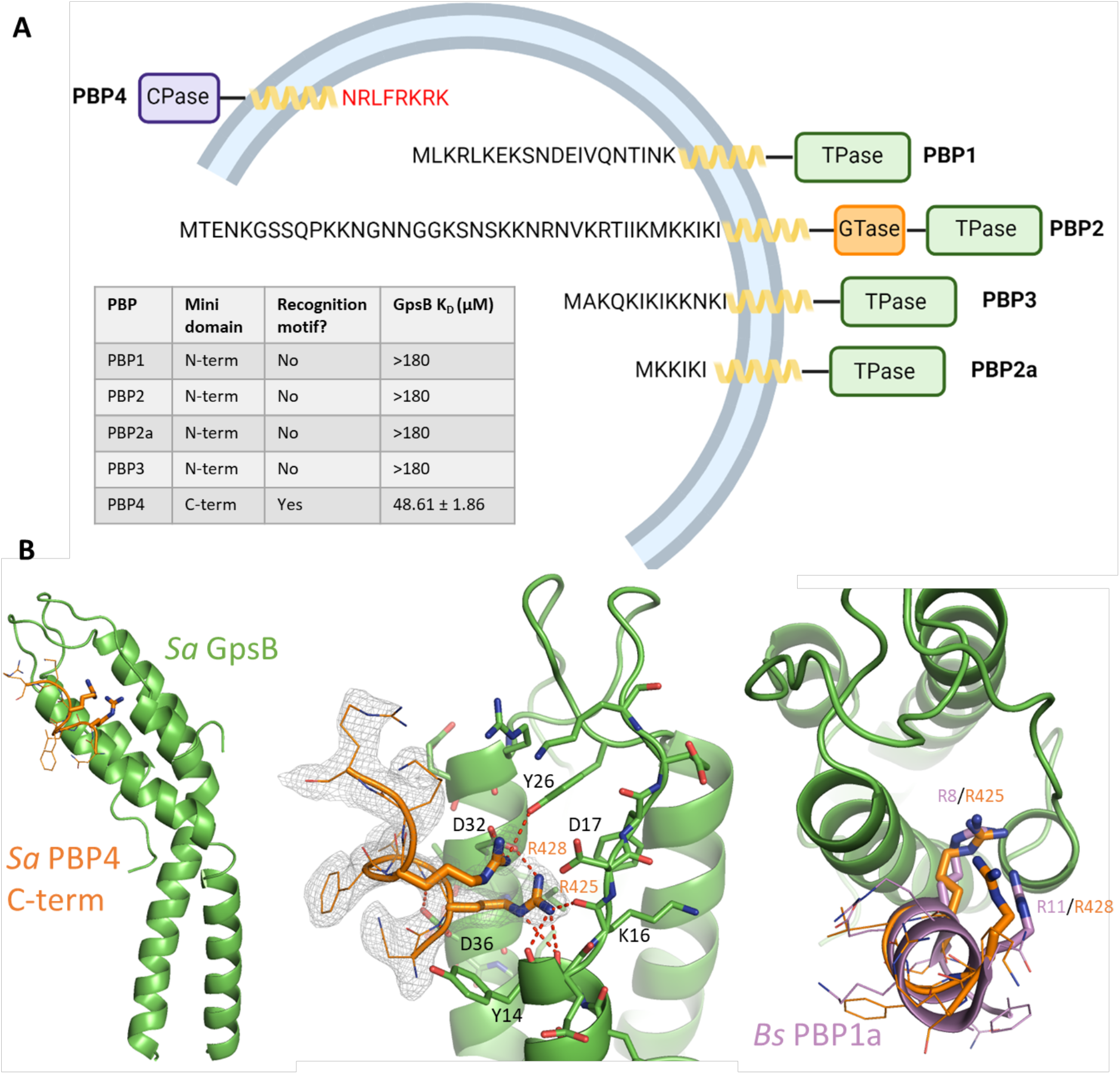
The C-terminal mini-domain of PBP4 directly interacts with GpsB. **(A)** Domain representation of the four/five *S. aureus* (COL)/MRSA (USA300) PBPs. Each protein is shown from the N-terminus (left) to the C-terminus (right). All four transpeptidase PBPs: PBP1, PBP2, PBP3, and PBP2a lack a GpsB-recognition motif on their N-terminal, cytosolic mini-domain. In contrast, the C-terminal mini-domain of PBP4, the sole *S. aureus* class C PBP, contains this motif (NRLFRKRK, red). The dissociation constants were determined with SPR (n=2). **(B)** Crystal structure of *Sa* GpsB R24A in complex with PBP4 C-terminal peptide fragment at 2.40 Å resolution. The middle panel includes the electron density map of the Sa PBP4 heptapeptide, 2F_o_-F_c_ = 1.0 σ. The right panel shows a superimposition of the *Bs* PBP1 mini-domain from the *Bs* GpsB + PBP1 complex (PDB ID 6GP7, purple) highlighting similar binding features.

Initial efforts to obtain a crystal structure of *Sa* PBP4 and *Sa* FtsZ with *Sa* GpsB were unsuccessful. A key factor preventing the formation of this complex were the tight interactions forming at the crystal packing interface. By extending the asymmetric unit, we found each GpsB dimer coordinates two others in a head-to-head arrangement **(Fig. S5 A, B)**. This involves the insertion of Arg and Lys residues from the membrane binding loop into the PBP binding site from the adjacent GpsB protomer, mimicking the binding mode observed between PBP mini-domains and GpsB in orthologous structures, such as *Bs* PBP1 (**Fig. S5 C,D**)^16^. It is unclear whether this head-to-head interaction occurs in the cell or is simply a crystallization artifact. Nonetheless, it is apparent this interaction would need to be disrupted to capture interactions with a binding partner. To do so, we generated an R24A mutant because of its central role in the head-to-head interaction and its distance from the PBP-binding groove. This point-mutation successfully disrupted the occluded crystal packing interface and allowed us to determine a 2.40 Å resolution structure of *Sa* PBP4 peptide bound to *Sa* GpsB (**Fig. 4B, Table S1**). Unambiguous electron density for the *Sa* PBP4 C-terminal octapeptide, corresponding to an α-helix, was resolved at the PBP-binding groove of GpsB. This structure clearly shows that two of these residues, Arg425 and Arg428, are key components of the *Sa* PBP4 - *Sa* GpsB interaction, where they form multiple hydrogen bonds with the main chain amides of *Sa* GpsB Ile13, Tyr14, Lys16 and the sidechain hydroxyl of Tyr26. Furthermore, Arg425 and Arg428 form two salt bridges with the carboxyl sidechain of Asp32. Additionally, *Sa* GpsB Asp36 appears to play a major role in stabilizing the PBP4 α-helix by forming two hydrogen bonds with the backbone nitrogen of Arg425 and Leu426, an arrangement that is only possible when these two residues are part of an α-helix. Overall, the complex of *Sa* PBP4/*Sa* GpsB is very similar to *Bs* PBP1/*Bs* GpsB (**Fig. S1, 4B**) and is distinguished by the interactions of two Arg residues with the backbone and acidic sidechains lining the PBP binding groove.

### *Sa* FtsZ and *Sa* PBP4 have lower affinity for *Sa* GpsB^ΔMAD^ _FL_ compared to *Sa* GpsB^WT^ _FL_

Due to the apparent importance of the three-residue insertion at the midpoint of *Sa* GpsB, we also tested the affinity of *Sa* FtsZ and *Sa* PBP4 derived peptides against *Sa* GpsB^ΔMAD^ _FL_. Using SPR we found that the affinity was significantly reduced for *Sa* GpsB^ΔMAD^ _FL_ compared to *Sa* GpsB^WT^ _FL_ (**Table S2**). Dose-dependent saturation was very weak (K_D_ > 200 μM) for *Sa* PBP4 (423-431) and *Sa* FtsZ (379-390), which verged on being undetectable. Additionally, titration of ^15^N-labelled *Sa* GpsB^ΔMAD^_1-70_ with the *Sa* FtsZ_383-390_ peptide does not result in chemical shift changes to the GpsB^ΔMAD^ ^1^H-^15^N HSQC, but only in broadening of a small number of signals, which is consistent with the higher K_D_ measured by SPR (**Fig. S2C**). While these mutants demonstrate better thermal stability (**Fig. 2A**), it is possible their deletion may disrupt interactions with α-helix 1 (res. 10-16; **Fig. 1B**), which may subsequently alter the size or shape of the PBP-binding pocket, thus reducing the favorable interactions that promote binding.

## Discussion

GpsB is an important protein that coordinates multiple elements of cell wall synthesis machinery. Although GpsB is widespread amongst Firmicutes, it has unique structural characteristics and functional roles in *S. aureus*. In this study, we initially present the crystal structure of the *Sa* GpsB N-terminal domain (1-70) (**Fig. 1A**). The characteristic coiled-coil motif demonstrates conformational flexibility resulting from a three-residue hinge region that is unique to *Staphylococci* (**Fig. 1 C, D, E**). The function of this hinge region and the flexibility it imparts remains unclear, but its deletion increases thermal stability (**Fig. 2A**) and weakens affinity for *Sa* FtsZ and *Sa* PBP4 (**Table S2**). Furthermore, unlike WT *Sa* GpsB, ΔMAD and ΔMNN mutants are not toxic to *B. subtilis* (**Fig. 2 C, D**), underscoring the cellular significance of this region. In *S. aureus*, we show that the phenotypes of ΔMAD (but not ΔMNN) closely resembles that of *Sa* GpsB (**Fig. S4A, C**). We posit that the difference is likely due to the higher affinity of ΔMNN to itself than *Sa* GpsB based on bacterial two-hybrid analysis (**Fig. S4B**). Regardless, both ΔMAD and ΔMNN mutants localize to the division site in both *S. aureus* and *B. subtilis* suggesting they retain some level of affinity for their usual interaction partners.

Next, we identified a GpsB-recognition motif on the C-terminus of both *Sa* FtsZ and Sa PBP4, leading to biophysical, biochemical, and structural experiments providing evidence of a direct interaction between these proteins and the N-terminal domain of *Sa* GpsB (**Fig. 3**). This recognition motif, which is also present in *Lm* PBPA1, *Sp* PBP2a, and *Bs* PBP1, involves the insertion of several Arg residues into the binding groove formed at the coiled-coil interface near the membrane binding loop (**Fig. S1**)^16^. Our group previously found that *Sa* GpsB interacts with *Sa* FtsZ to regulate its polymerization characteristics, but the repeated (S/T/N)-R-X-X-R-(R/K) GpsB recognition motif on its C-terminus was not recognized until the work on such motifs in PBPs published by Cleverley *et al.* 2019^16^. This motif is unique to *Staphylococci* FtsZ and is absent/less conserved in other Firmicutes **(Fig. 3A)**. It is unclear why the GpsB recognition motif is repeated, since there is only enough area in the PBP-binding groove to accommodate a helix of eight to nine residues. It is possible that *Sa* FtsZ binds two *Sa* GpsB dimers simultaneously, given the putative arrangement of GpsB as trimer of dimers^29^. Alternatively, the first site may be occluded by the binding of other FtsZ interaction partners that are known to bind FtsZ through the C-term such as FtsA, EzrA, and SepF^36^. This finding, in addition to our previous reports^12,15^, underscore the importance of GpsB in *S. aureus* cell division.

A crystal structure of the *Sa* PBP4 C-terminal recognition sequence bound to the N-terminal domain of GpsB, reveals a binding mode that mimics previously solved orthologous PBP/GpsB pairs. This discovery is notable because it was previously thought that only the N-terminal “mini-domain” of transpeptidase PBPs (class A, class B) could bind to GpsB. Furthermore, this finding is significant not only because it is the sole *S. aureus* PBP that binds to GpsB, but also because *Sa* PBP4 is intimately associated with WTA synthesis^23,37^. Previous studies have found that *Sa* GpsB interacts with the WTA biosynthesis pathway proteins TarO^14^ and TarG^15^, and likely facilitates the export of WTA to the septal cell wall. To our knowledge, *Sa* PBP4 does not directly interact with any of the known WTA machinery. However, it does require WTA for recruitment to the division septum, and impairment of WTA assembly results in delocalization of *Sa* PBP4^23^. Thus, while *Sa* PBP4 itself is not essential for growth, it is tightly regulated by WTA synthesis, an essential process that is also mediated by GpsB, which associates with WTA proteins TarO and TarG^14,15^. Thus, it is conceivable that GpsB likely arrives at the division site together with FtsZ, facilitates WTA synthesis, and subsequently recruits PBP4 to promote efficient cytokinesis. PBP4 joins several other known proteins found to interact with *Sa* GpsB: EzrA^13^, DivIVA^38^, FtsZ^12^, TarO^14^, and TarG^15^ **(Fig. 5)**. The interaction diagram outlines the known interactions for GpsB to date but given the diverse number of proteins at the divisome and various binding surfaces on GpsB, it will surely be expanded in the future. Furthermore, while GpsB lacks certain direct interactions with other known divisome proteins, they are indirectly linked through intermediate proteins. For example, *Sa* FtsZ binds to *Sa* EzrA, a protein known to interact with the SEDS–PBP pairs *Sa* PBP1-FtsW and *Sa* PBP3-RodA, thus indirectly coupling *Sa* GpsB to enzymes that are critical for peptidoglycan synthesis at the divisome^13,39^.

**Figure 5.**
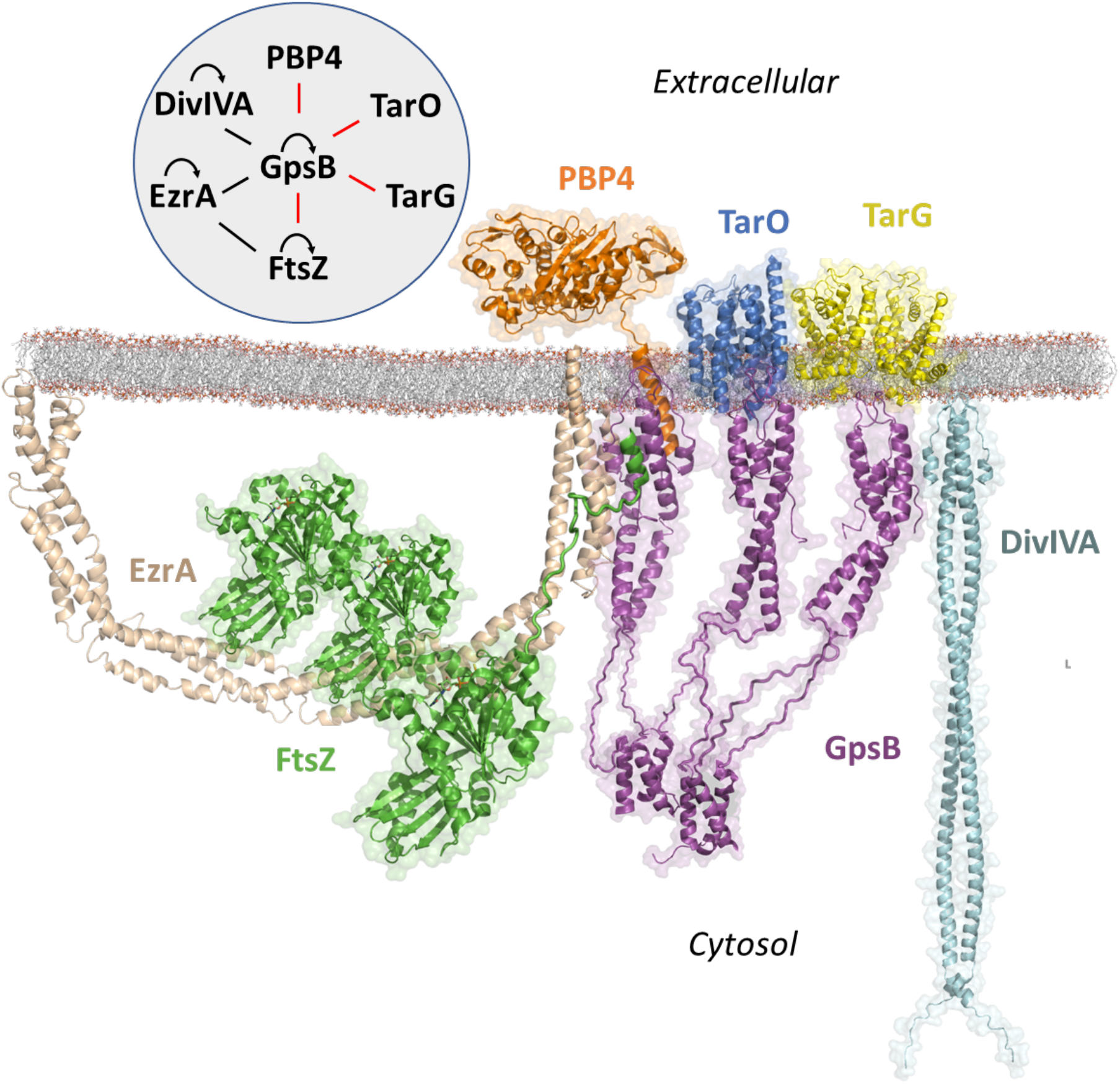
Known interactome of *S. aureus* GpsB and putative arrangement at the division septum in graphical and diagram (upper left) format. In this paper we demonstrate that the C-terminal mini-domain of PBP4 (orange) and the C-terminus of FtsZ (green) bind to the N-terminal domain of GpsB (purple). The regions within GpsB responsible for interacting with other partners remain to be elucidated. For interaction diagram, the red lines indicate interactions putatively unique to *S. aureus*, black lines indicate interactions found in both *S. aureus* and *B. subtilis*, and curved arrows represent self-interaction.

When evaluating the biophysical affinity of *Sa* FtsZ and *Sa* PBP4 for *Sa* GpsB, it is important to consider the peptide dissociation constants determined by SPR (~ 20 to 80 μM) may not directly translate to, and likely underestimate, the cellular affinity. The divisome is a highly complex environment with multiple proteins that interact near or at the cell membrane. The enrichment of both *Sa* FtsZ and *Sa* PBP4 at the division septum likely increases their apparent affinity simply based on avidity. Additionally, the arrangement of proteins may introduce synergistic interactions. Studies have found that EzrA, which binds to *Sa* GpsB^13^, also binds to the C-terminus of FtsZ^34,40^, likely upstream of the *Sa* GpsB recognition motif in *S. aureus*. This interaction could both increase the local concentration of *Sa* FtsZ and induce molecular recognition features that promote association with *Sa* GpsB. In addition, the oligomeric state of FtsZ and GpsB may further lead to cooperativity in the interactions between the FtsZ filament and GpsB hexamers. However, tighter binding may also prove deleterious in certain scenarios, especially for dynamic proteins like FtsZ. Given that *Sa* GpsB enhances the GTPase activity (required for FtsZ filament disassembly) of *Sa* FtsZ^12^, it is possible GpsB could dynamically promote polymerization and depolymerization of FtsZ.

Under the specter of growing antibacterial resistance, the identification of novel antibiotic targets is becoming increasingly urgent. Beyond PBPs, the bacterial divisome is a largely untapped source of antibiotic targets. The delineation of the physiological role of *Sa* GpsB, identification of its recognition motifs, and characterization of its 3D structure greatly enables modern antibiotic drug-discovery strategies. Furthermore, the involvement of *Sa* GpsB in multiple essential processes presents the opportunity to design targeted antibiotics. There is a growing need for narrow-spectrum antibiotics, especially for common infections, such as those caused by *S. aureus*^41^. Narrow spectrum antibiotics avoid selective pressure of commensal bacteria which can serve as a reservoir for resistance elements. In the same vein, their limited disruptive properties can avoid pathologies associated with bacterial dysbiosis, like *Clostridioides difficile* infection. The results from this study provide new information for both understanding the role of GpsB in *S. aureus* division and probing new avenues for narrow-spectrum antibiotic development.

## Methods

### Recombinant protein cloning and purification

The nucleotide sequence of *S. aureus gpsB* corresponding to the N-terminal domain (1-70) and full length (1-114) was inserted into a modified pET28a vector with a His-TEV sequence 5’ to the multiple cloning site (MCS). GpsB constructs expressed for SPR bioanalysis were cloned into a separate pET28a vector with a His-TEV-Avi sequence flanking the MCS (pAViBir). ΔMAD and ΔMNN mutants were generated with QuikChange site directed mutagenesis using custom primers (ΔMAD (5’- GATTATCAAAAAATGAATAATGAAGTTGTAAAATTATCAGAAGAGAATC) and ΔMNN (5’- GATTATCAAAAAATGGCCGATGAAGTTGTAAAATTATCAGAAGAG)). All plasmids were transformed into Rosetta (DE3) pLysS cells. A single colony was grown in LB media supplemented with 35 μg/mL chloramphenicol and 50 μg/mL kanamycin at 37 °C overnight. The overnight culture was then diluted into 1 L media at 1:500 and incubated at 37 °C until the OD_600_ reached 0.8. Protein expression was initiated with 0.5 mM IPTG and continued incubation at 25 °C overnight. pAviBir contructs were biotinylated during IPTG induction with a stock of 5 mM biotin dissolved in bicine for a final concentration of 50 μM. Cells were harvested by centrifugation at 5,000 x *g* for 10 min. The cell pellet was resuspended in buffer A (20 mM Tris-HCl pH 8.0, 300 mM NaCl, 20 mM imidazole, and 10 % glycerol). Cells were disrupted by sonication followed by centrifugation at 35,000 x *g* for 40 min. The pellet containing the protein was resuspended in buffer AD (100 mM Tris-HCl pH 8.0, 6 M guanidine HCl, 300 mM NaCl) and incubated at 30 °C for ~one hour to fully dissolve the pellet, followed by centrifugation at 45,000 x *g* for one hour. The supernatant was then loaded onto a HisTrap affinity column and eluted in a single step using Buffer BD (100 mM sodium acetate pH 4.5, 6 M guanidine HCl, 300 mM NaCl). The eluted protein was diluted dropwise in refolding buffer (100 mM Tris pH 8.0, 200 mM NaCl) and allowed to refold overnight at 4 °C. The sample was then loaded to a HisTrap column and eluted with liner gradient of imidazole. The fractions containing GpsB were pooled and concentrated. The protein was then incubated with TEV at 1:20 ratio overnight at 4 °C. The sample was then loaded onto a HisTrap for reverse Ni^2+^ cleanup. Flow through was collected and purified using a HiLoad 16/60 Superdex 75 size exclusion column. The protein was stored at −80 °C in the storage buffer (20 mM Tris-HCl pH 8.0, 200 mM NaCl). The purity of the protein was determined by SDS-PAGE as >95%.^15^N-labelled GpsB (1-70) for NMR spectroscopy was expressed in the same way, but in minimal media containing ^15^N NH_4_Cl and ^12^C glucose as the source of nitrogen and carbon, respectively, supplemented with MgSO_4_, CaCl_2_, and trace metals.

### Strain construction

Plasmids were generated using standard cloning procedures. The C-terminal six-residue truncation of FtsZ (FtsZ ^ΔC6^) was generated by PCR using primer pairs oDB9/oDB10 (NdeI/XhoI) and cloned into the pET28a to create pDB1. FtsZ CTT encoding the C-terminal 66 amino acids of *S. aureus* FtsZ was cloned into pET28a using oP228/oP229 (Nde1/BamHI) to create pSK4. These were then transformed into BL21-DE3 cells creating EDB01 and SK7 respectively. ΔMAD was created by using site directed mutagenesis with custom primers (5’-GATTATCAAAAAATGAATAATGAAGTTGTAAAATTATCAGAAGAGAATC) and ΔMNN was ordered from Integrated DNA Technologies as oLHgblock1. These mutations were then PCR amplified and cloned into pDR111 to create both untagged (oP36/oP38; Hind111/Sph1) and GFP tagged (oP36/oP37; Hind111/Nhe1 and oP46/oP24; Nhe1/Sph1) variants. Plasmids were then transformed into PY79 and screened for amyE integration resulting in strains LH115, LH116, LH119, and LH126. The PCR products containing the ΔMAD and ΔMNN mutations were also cloned into pCL15 backbone using the same primers and restriction sites to create plasmids, pLH62, pLH59, pLH63, and pLH60. These plasmids were then transformed into RN4220 cells resulting in strains LH129, LH127, LH130, and LH128. Then these plasmids were transduced into SH1000 to create strains LH135, LH134, LH133, and LH132. To make the BTH plasmids, ΔMAD and ΔMNN were amplified (BTH11/BTH12; EcoRI/XhoI) and cloned into pEB354 and pEB355 resulting in strains LH164, MA1, LH170, and LH168. The genotypes of strains and oligonucleotides used in this study are provided in **Tables S3** and **S4**.

### X-ray crystallography

Crystals of GpsB (1-70) were grown in a hanging drop apparatus by mixing 11 mg/mL GpsB (purity > 98%) with crystallization buffer (30% PEG 3350, 0.4 M NaCl, and 0.1 M Tris pH 8.5) in an equal ratio at 20 °C. Filamentous crystals appeared overnight and were harvested after one week of growth by briefly transferring to cryoprotectant (30% PEG 3350, 0.4 M NaCl, 0.1 M Tris pH 8.5, and 15% glycerol), followed by flash freezing in liquid nitrogen. GpsB (1-70) and PBP4 (424-451) were mixed in a 1:1 ratio with a GpsB concentration of 6 mg/mL and 1.44 mM PBP4 peptide (1:1 ratio). Initial crystals grew in 25% PEG 4000, 0.1 M Tris pH 8.0, and 0.2 M sodium acetate. These crystals were crushed, diluted 10,000-fold, then seeded into drops in a ratio of 1:1:0.5 (protein, crystallization solution, seed stock). Crystals were harvested after one week of growth by transferring to a cryoprotectant solution of 27.5% PEG 4000, 0.1 M Tris pH 8.0, 0.2 M sodium acetate, and 15% glycerol. X-ray diffraction data were collected on the Structural Biology Center (SBC) 19-ID beamline at the Advanced Photon Source (APS) in Argonne, IL, and processed and scaled with the CCP4 versions of iMosflm^42^ and Aimless^43^. Initial models were obtained using the MoRDa^44^ package of the online CCP4 suite. Unmodeled regions were manually built and refined with Coot^45^.

### Circular dichroism

Thermal stability was assessed with circular dichroism using a Jasco J-815 CD spectropolarimeter coupled to a Peltier cell holder. Recombinantly expressed and purified WT GpsB and GpsB mutants were diluted to 2 μg/ml in 50 mM sodium phosphate (pH 7.0) and CD spectra were measured at 222 nm from 20 °C to 80 °C. Melting temperature was determined with a 4-parameter logistic curve fit using Graphpad Prism 9.

### Surface plasmon resonance

A Series S CM5 chip (Cytiva) was docked into a Biacore S200 instrument (Cytiva) followed by surface activation with NHS/EDC amine coupling. Lyophilized neutravidin (Thermo Fisher Scientific) was dissolved in sodium acetate pH 5.25 to a final concentration of 0.25 mg/mL and injected onto the activated CM5 chip at 10 μL/min for 5 min. Biotinylated GpsB was diluted to 1 mg/mL in HEPES-buffered saline (HBS) and injected over the neutravidin-immobilized CM5 chip at 20 μL/min for 5 min. Synthetic peptides and recombinantly expressed *Sa* FtsZ (325-390) were serially diluted in HBS, and injected at 30 μL/min for 50 s with a dissociation of 100 s, followed by a stabilization period of 15 s and a buffer wash between injections. All experiments were performed in technical duplicate. Dissociation constants were determined with a one site binding model using Graphpad Prism 9.

### NMR spectroscopy

2D ^1^H-^15^N HSQC spectra of *Sa* GpsB (1-70) were recorded on an Agilent 800-MHz direct drive instrument equipped with a cryoprobe. NMRpipe^46^ and Sparky (University of California, San Francisco) were used for processing and analysis, respectively. All spectra were acquired in 20 mM Tris pH 8.0, 200 mM NaCl, prepared in 7.5% D_2_O, and at 25 °C. The concentration of *Sa* GpsB^WT^_1-70_ was 160 or 700 μM for the free protein spectrum and 160 μM for the complex with the *Sa* FtsZ octapeptide, which was added in a 1.5x molar excess.

### GTP hydrolysis of FtsZ

*Sa* FtsZ, *Sa* GpsB, and *Sa* FtsZ^ΔC6^ were purified using Ni-NTA affinity chromatography as described previously^12^. The effect of GpsB on the GTPase activity of FtsZ and FtsZ^ΔC6^ was determined by measuring the free phosphate released using the malachite green phosphate assay kit (Sigma Aldrich). Briefly, either FtsZ or FtsZ^ΔC6^ (30 μM) was incubated with GpsB (10 μM) in the polymerization buffer (20 mM HEPES pH 7.5, 140 mM KCl, 5 mM MgCl_2_) containing 2 mM GTP at 37 °C for 15 min. The free phosphate released was determined by measuring the absorbance of the reaction mixture at 620 nm.

### *B. subtilis* and *S. aureus* growth conditions

Liquid cultures of *B. subtilis* cells were grown in LB and *S. aureus* cells were grown in TSB supplemented with 10 μg/mL chloramphenicol at 37 °C.

### Spot titer assays

Overnight cultures of *B. subtilis* and *S. aureus* strains were back diluted to OD_600_ = 0.1 and grown to midlog phase (OD_600_ = 0.4). Cultures were then back diluted to an OD_600_ of 0.1, serial diluted, and spotted onto LB plates with or without 1 mM IPTG (*B. subtilis*), or TSA plates supplemented with 10 μg/mL chloramphenicol with or without 1 mM IPTG (*S. aureus*), and incubated overnight at 37 °C.

### Fluorescence microscopy

Overnight cultures of *B. subtilis* and *S. aureus* strains to be imaged were back diluted to OD_600_ = 0.1 and grown to midlog phase (OD_600_ = 0.4) and then induced with 1 mM IPTG and allowed to grow for an additional 2 h. Cells were then prepared and imaged as previously described^47^. Briefly, 1 mL aliquots were spun down, washed, and resuspended in PBS. Cells were then stained with 1 μg/mL SynaptoRed fluorescent dye (Millipore-Sigma) to visualize the membrane and 5 μL of culture was spotted onto a glass bottom dish (Mattek). Images were captured on a DeltaVision Core microscope system (Leica Microsystems) equipped with a Photometrics CoolSnap HQ2 camera and an environmental chamber. Seventeen planes were acquired every 200 nm and the data were deconvolved using SoftWorx software. Cells were measured using ImageJ and analyzed using GraphPad Prism 9.

### Bacterial two-hybrid assay

Plasmids carrying genes of interest cloned into the pEB354 (T18 subunit) and pEB355 (T25 subunit) backbones were transformed pairwise into BTH101 cells. Overnight cultures of the strains grown in 100 μg/mL ampicillin, 50 μg/mL kanamycin, and 0.5 mM IPTG, at 30 °C, were spotted onto McConkey agar containing 1% maltose that were also supplemented with ampicillin, kanamycin, and IPTG. Plates were incubated for 24 h at 30 °C and then imaged. The β-galactosidase assay was carried out as previously described^15^. Mixtures of 20 μL of culture, 30 μL of LB, 150 μL Z buffer, 40 μL ONPG (4 mg/mL), 1.9 μL β-mercaptoethanol, and 95 μL polymyxin B (20 mg/mL), were transferred to a 96-well plate and read on a BioTek plate reader. Miller units were then calculated and graphed using GraphPad Prism 9.

### Immunoblot

Overnight cultures of *S. aureus* cells were back diluted to OD_600_ = 0.1, grown to midlog phase (OD_600_ = 0.4), and then induced with 1 mM IPTG and grown for an additional 2 h. Cells were then standardized to an OD_600_ = 1.0, lysed with 5 μL lysostaphin, and incubated for 30 min at 37 °C. 1 μL of DNAse A (1 U/μL) was added and incubated for an additional 30 min. Samples were then analyzed by SDS-PAGE analysis, transferred to a membrane, and probed with rabbit antiserum raised against GpsB-GFP. Total protein was visualized from the SDS-PAGE gel using the GelCode Blue Safe Protein Stain (ThermoFisher).

### Homology model construction of deletion variants

Template-based homology models were made using MODELLER 9.24^48^ by constructing 1,000 decoys corresponding to each construct based on various template structures.

### Molecular dynamics (MD) simulations

The GpsB dimers were exposed to conventional MD simulation using Gromacs (v. 5.0.4)^49,50^ with the CHARMM36m force field^51^. Explicit TIP3P water^52^ with 150 mM KCl was used for solvation. A 12 Å cut-off for the van der Waals forces was used. Electrostatic forces were computed using the particle mesh Ewald method^53^. The Verlet cut-off scheme was used. The temperature and pressure were controlled using the Nosé-Hoover^54–56^ and Parrinello-Rahman^57,58^ methods, respectively, to sample the NPT ensemble at P = 1 bar and T = 303.15 K. The integration time step was 2 fs, enabled by using H-bond restraints^59^. Each system was simulated for 250-500 ns. All systems were made using CHARMM-GUI^60–62^.

## Data Availability

All crystal structures have been deposited in the RCSB Protein Data Bank (PDB) with accession IDs of: *Sa* GpsB NTD (PDB ID 8E2B), *Sa* GpsB NTD + *Sa* PBP4 C-term (PDB ID 8E2C).

## Acknowledgements

We thank Eric Lewandowski for reading the manuscript. We also thank the staff members of the Advanced Photon Source of Argonne National Laboratory, particularly those at the Structural Biology Center (SBC) for assistance with X-ray diffraction data collection. SBC-CAT is operated by UChicago Argonne LLC, for the U.S. Department of Energy, Office of Biological and Environmental Research under contract DE-AC02-06CH11357.

## Funding

This work was supported by the NIH (R21 AI164775 (Y.C. and P.J.E.) and R35 GM133617 (P.J.E.)).

## Author contributions

The studies presented herein were conceived and designed by M.D.S., L.R.H., P.J.E., and Y.C.; The manuscript was written by M.D.S., L.R.H., I.G., P.J.E., and Y.C.; Y.C., P.J.E., and I.G. provided scientific input, funded and supervised the studies; Figures were prepared by M.D.S., L.R.H., and I.G.; Protein constructs were designed by M.D.S. and Y.C.; Proteins were cloned and purified by X.Z., S.J.K., and D.B.; Surface plasmon resonance bioanalysis was performed by M.D.S.; Thermal shift binding assays were performed by M.T.K.; Crystallization experiments were performed by M.D.S. with help from S.G.B. and A.C.J.; Crystal structures were solved and refined by M.D.S.; I.G. and R.E.N. performed NMR analysis; GTPase assay was performed by D.B.; Fluorescence microscopy, bacterial two hybrid analysis, and growth assays were conducted by L.R.H.; Protein modeling and MD simulations were performed by J.J.M.

## Competing interests

The authors declare no competing interests.

## Supplementary Data

**Supplementary Figure 1.**
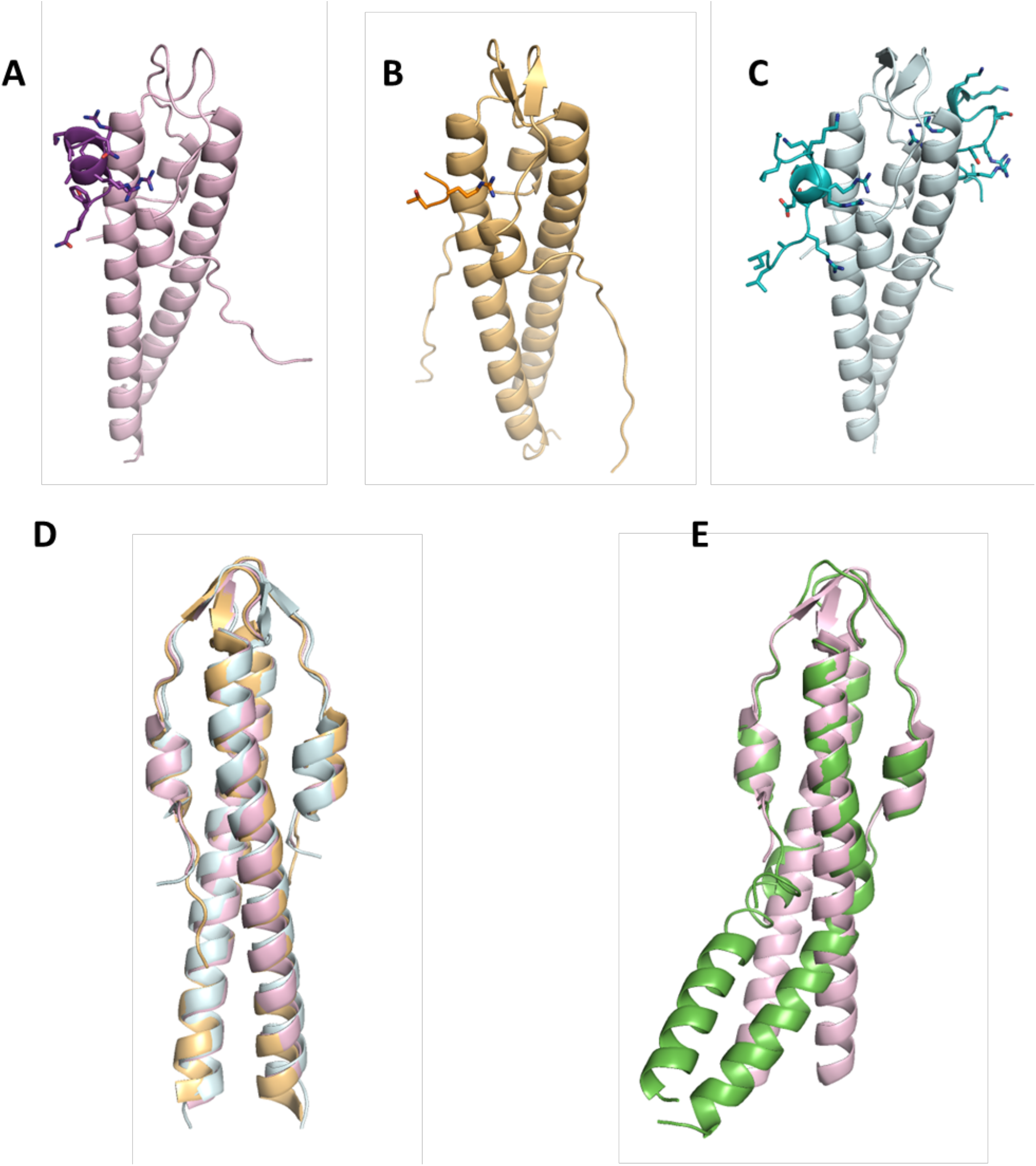
Crystal structures of PBP mini-domains in complex with the N-terminal domain of their cognate GpsB, published by Cleverley *et al.* 2019^16^. **(A)** *Bs* PBP1. **(B)** *Lm* PBPA1 **(C)** *Sp* PBP2a **(D)** Superimposition of previously solved GpsB structures from subpanels A-C. **(E)** Superimposition of *Sa* GpsB with *Bs* GpsB from subpanel A.

**Supplementary Figure 2.**
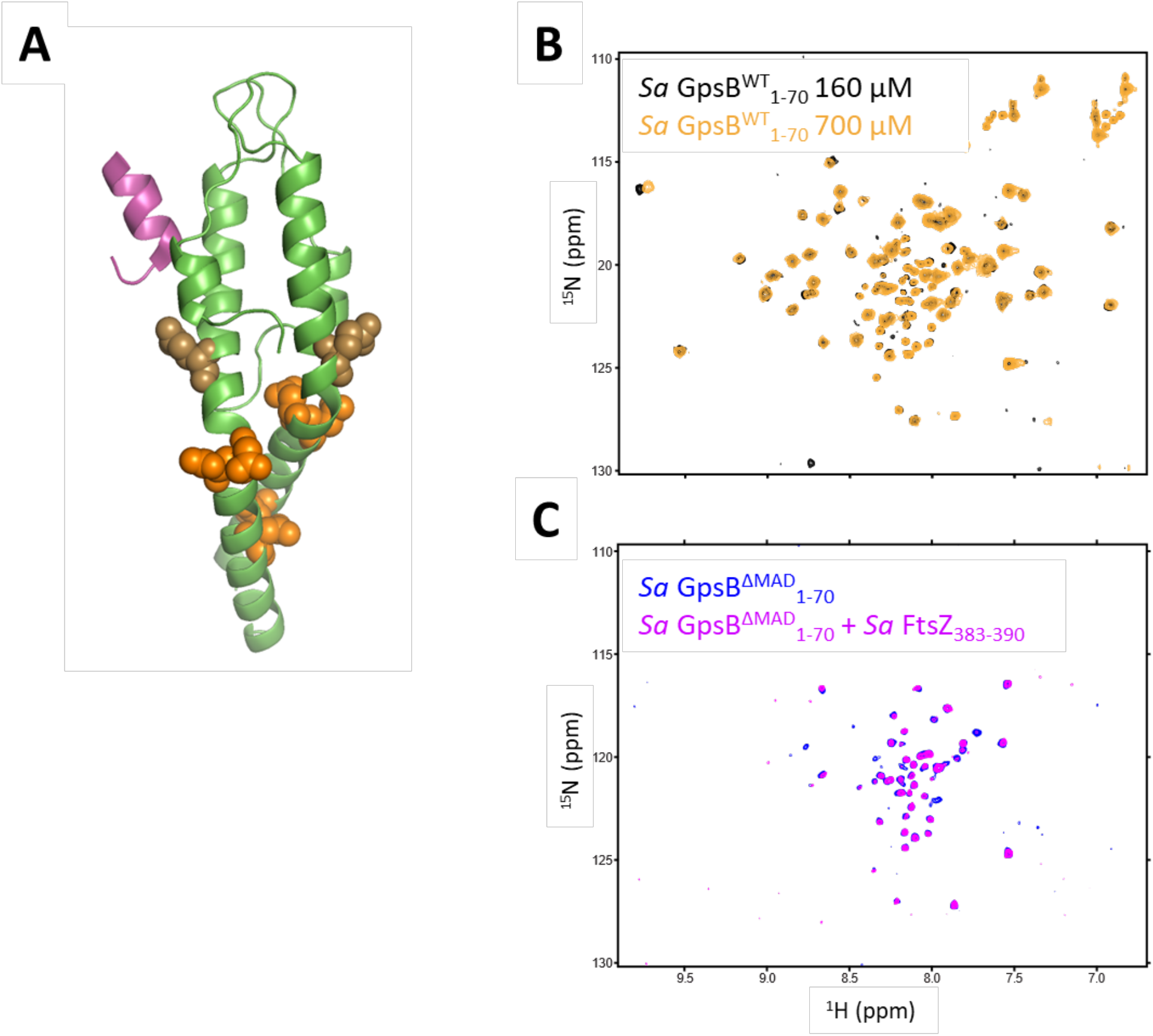
**(A)** Asn (orange) and Gln (brown) residues mapped on the structure of GpsB, modeled with a PBP-derived peptide (purple) from *Bs* PBP1A (PDB ID 6GP7). **(B)** Overlays of the ^1^H-^15^N HSQC of spectrum of *Sa* GpsB^WT^_1-70_ acquired at different concentrations. **(C)** The ^1^H-^15^N HSQC of *Sa* GpsB^ΔMAD^_1-70_ in the absence (blue) and presence of 1.5 equivalents of *Sa* FtsZ_383-390_ peptide (magenta).

**Supplementary Figure 3.**
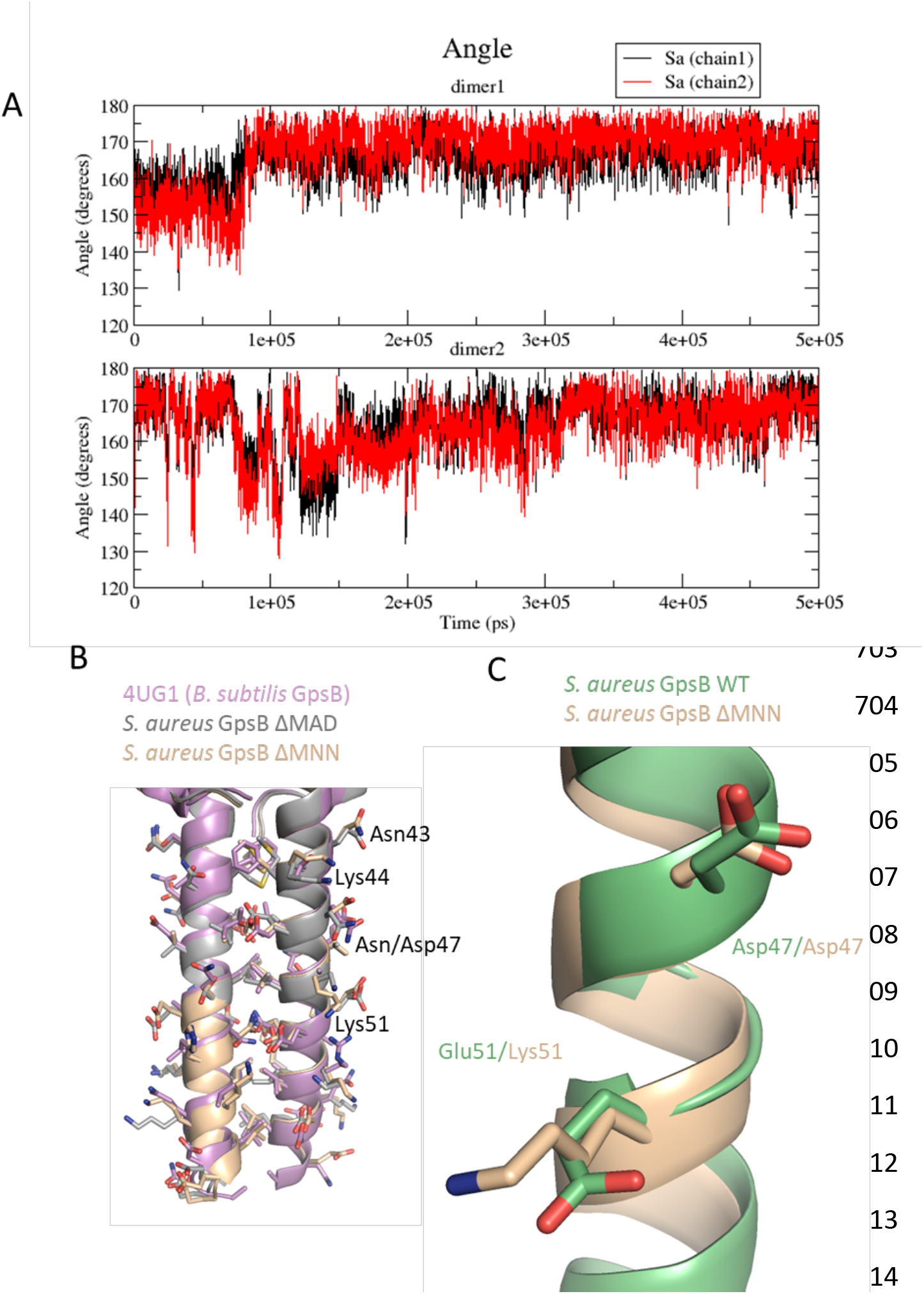
**(A)** Molecular dynamics simulation of the WT GpsB N-terminal domain. Significant fluctuations in pitch angle are observed within 100 ns for both dimers from the crystal structure that underwent separate simulations, and for even longer periods of time for dimer 2. **(B)** Superimposed homology model of *S. aureus* GpsB ΔMNN (tan), *S. aureus* GpsB ΔMAD (dark grey), and the crystal structure of *B. subtilis* GpsB (purple; PDB ID 4UG1). Residues following the MNN or MAD deletion are depicted, which are more consistent with the multiple sequence alignment presented in **Fig. 1E**. PDB coordinates are provided in source data. **(C)** Several residues form preferential interactions following the MAD and MNN deletions. Shown here is an example of a potentially new electrostatic interaction between Asp47 and Lys51 in ΔMNN mutant. This +/− pair replaces the unfavorable −/− Asp/Glu pair found in *S. aureus* GpsB WT. Asp47 is replaced by Asn47 in ΔMAD mutant.

**Supplementary Figure 4.**
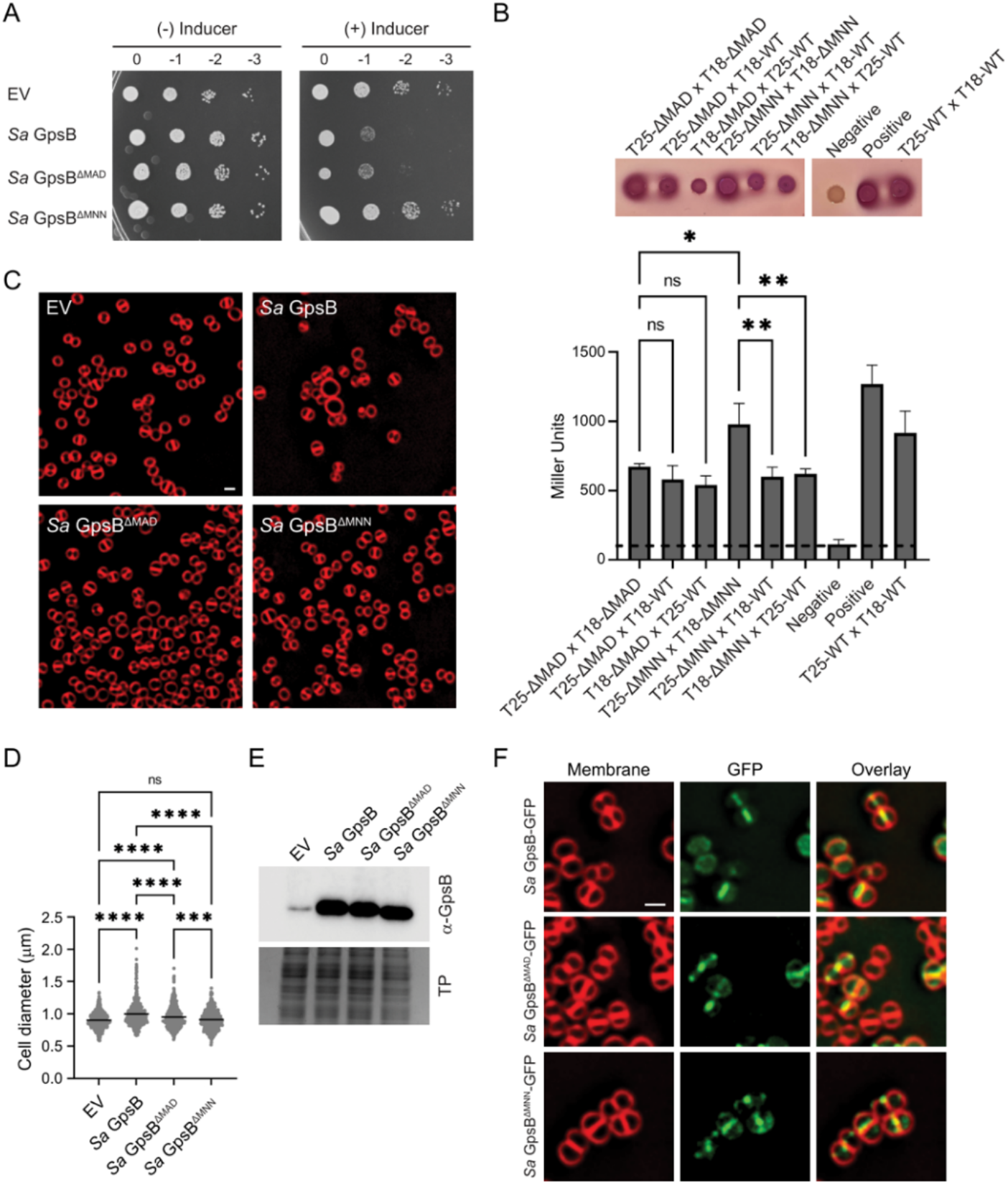
**(A)** Growth assay: serial dilutions of *Sa* strains harboring inducible *Sa* GpsB (GGS2), *Sa* GpsB ΔMAD (LH129), *Sa* GpsB ΔMNN (LH127), and the EV control (PE355), plated on TSA plates supplemented with 10 μg/mL chloramphenicol both without (left) and with (right) 1 mM IPTG. **(B)** Bacterial two-hybrid analysis. Pairwise interactions of wildtype *Sa* GpsB (WT) (LH39/LH40) with the ΔMAD (LH164/MA1) and ΔMNN (LH170/LH168) mutants as well as the ΔMAD and ΔMNN self-interactions plated on McConkey agar plates supplemented with 1% maltose (top). Interactions were also tested by β-galactosidase assay (bottom). Assay was done in triplicate and the dashed line shows the average Miller Unit level of the negative control. * P < 0.05 and ** P < 0.01. **(C)** Micrographs of *S. aureus* cells containing plasmids with inducible *Sa* GpsB (PES13), *Sa* GpsB ΔMAD (LH135), and *Sa* GpsB ΔMNN (LH127), and the EV control (PES5), imaged 2 h post induction. Cells were visualized with 1 μg/mL SynaptoRed membrane dye. Scale bar is 1 μm. **(D)** Quantification of micrographs shown in panel C. n = 500 cells; *** P < 0.001 and **** P < 0.0001. **(E)** Western blot of strains shown in panel C. Note: The GpsB band in EV lane is of native GpsB. **(F)** Micrographs showing the localization of *Sa* GpsB-GFP (PES6), *Sa* GpsB ΔMAD (LH133), and *Sa* GpsB ΔMNN (LH132). Cells visualized with 1 μg/mL SynaptoRed Membrane Dye. Scale bar is 1 μm.

**Supplementary Figure 5.**
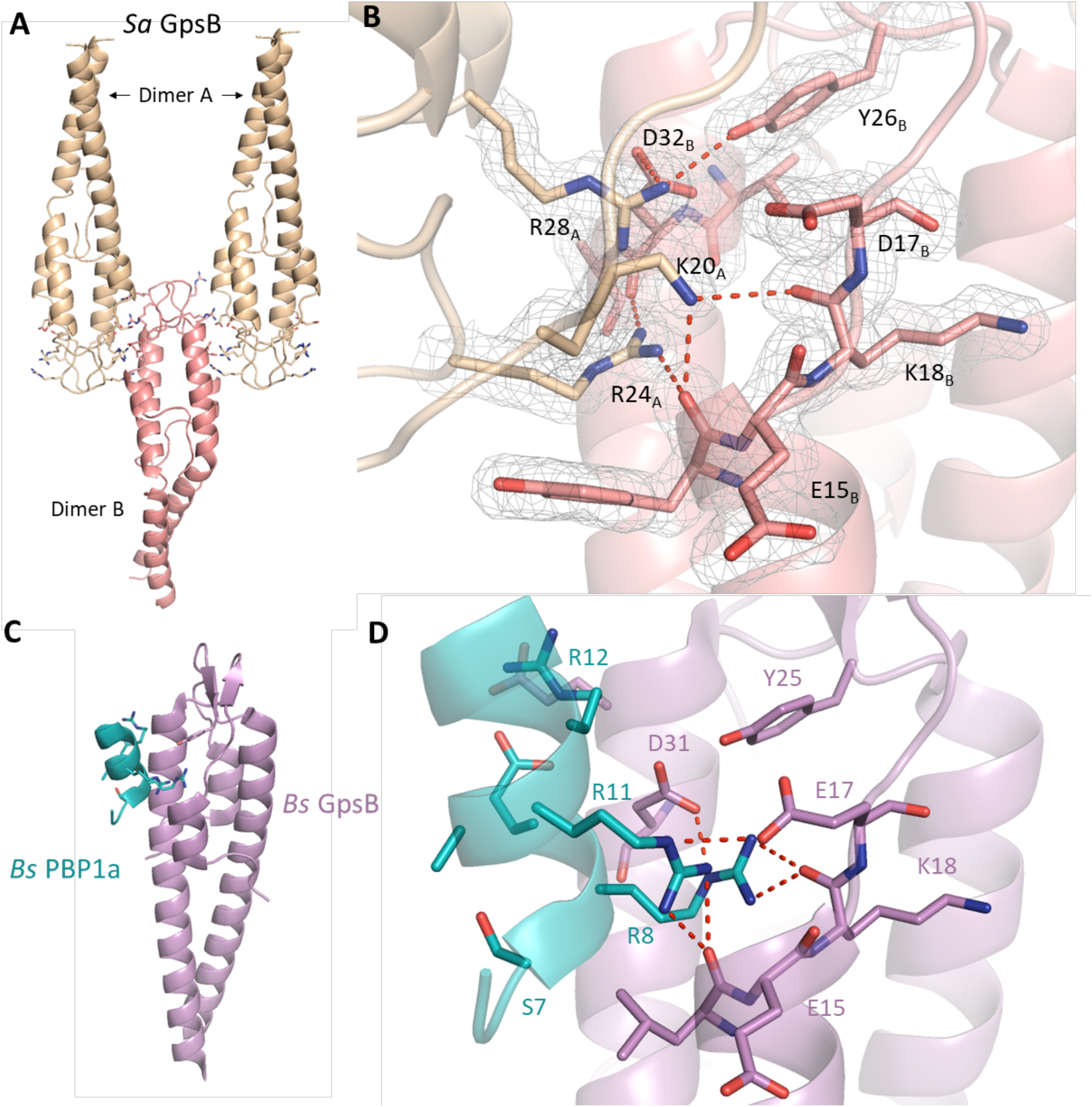
The crystal packing interface of *Sa* GpsB dimers form interactions that mimic those between GpsB-PBP pairs. **(A)** *Sa* GpsB dimers assemble in a head-to-head, antiparallel arrangement, where the membrane binding loop interacts with the PBP-binding groove of the other. **(B)** Enhanced viewpoint of the crystal packing interactions. 2F_o_-F_c_ map, shown in grey, is contoured at 1.0 σ. R28, R24, and K18 adopt similar interactions with the negatively charged PBP-binding groove, as the arginine fingers of **(C)**, **(D)** of *Bs* PBP1a with *Bs* GpsB (6GP7).

**Table S1.** Table of crystallographic statistics * Values in parentheses indicate those for the highest resolution shell.

| PDB ID | 8E2B | 8E2C |
| --- | --- | --- |
| Protein | Sa GpsB NTD | Sa GpsB NTD +<br>Sa PBP4 C-term |
| **Data Collection** |  |  |
| Space Group | P 1 2<sub>1</sub> 1 | P 1 2<sub>1</sub> 1 |
| Cell Dimensions |  |  |
| a, b, c (Å) | 53.52, 37.47,<br>79.25 | 29.35, 73.76,<br>42.37 |
| α, β, γ (°) | 90.00, 100.62,<br>90.00 | 90.00, 103.56,<br>90.00 |
| Resolution (Å) | 47.88 – 1.95 | 41.19 – 2.40 |
|  | (2.00 – 1.95) | (2.49 – 2.40) |
| R<sub>merge</sub> | 0.073 (0.190) | 0.095 (0.410) |
| <I>/σ<I> | 9.7 (4.1) | 10.8 (4.9) |
| Completeness (%) | 90.7 (94.8) | 95.7 (96.6) |
| Redundancy | 3.3 (3.2) | 5.2 (5.2) |
| **Refinement** |  |  |
| Resolution (Å) | 40.29 - 1.95 | 36.88 - 2.4 |
|  | (2.02 - 1.95) | (2.49 - 2.40) |
| No. reflections/free | 20762 / 1075 | 6579 / 633 |
| R<sub>work</sub>/R<sub>free</sub> | 0.188 / 0.240 | 0.227 / 0.248 |
| Clashscore | 4.92 | 5.63 |
| No. Atoms |  |  |
| Overall | 2768 | 1161 |
| Protein | 2335 | 1157 |
| Ligand/ion | 6 | 0 |
| Water | 427 | 4 |
| B-Factors (Å<sup>2</sup>) |  |  |
| Overall | 24.38 | 40.78 |
| Protein | 22.12 | 40.80 |
| Ligand/ion | 35.19 | - |
| Solvent | 36.55 | 33.34 |
| RMS Deviations |  |  |
| Bond Lengths (Å) | 0.015 | 0.013 |
| Bond Angles (°) | 1.68 | 1.70 |
| Ramachandran Favored (%) | 99.63 | 99.25 |
| Ramachandran Allowed (%) | 0.37 | 0.75 |
| Ramachandran Outliers (%) | 0.00 | 0.00 |
| Rotameric Outliers (%) | 3.14 | 2.40 |

**Table S2.** SPR dissociation constants (K_D_) of *Sa* FtsZ and *Sa* PBP4 derived peptides for *Sa* GpsB^WT^_FL_ and *Sa* GpsB^ΔMAD^_FL_

|  | *Sa* GpsB<sup>WT</sup><sub>FL</sub> | *Sa* GpsB<sup>ΔMAD</sup><sub>FL</sub> |
| --- | --- | --- |
| *Sa* FtsZ (325-390) | 40.21 ± 1.77 μM | 74.01 ± 4.34 μM |
| *Sa* FtsZ (379-390) | 73.63 ± 9.43 μM | >200 μM |
| *Sa* FtsZ (383-390) | 17.76 ± 1.25 μM | 59.68 ± 4.49 μM |
| *Sa* PBP4 (423-431) | 48.61 ± 1.38 μM | >200 μM |

**Table S3.**
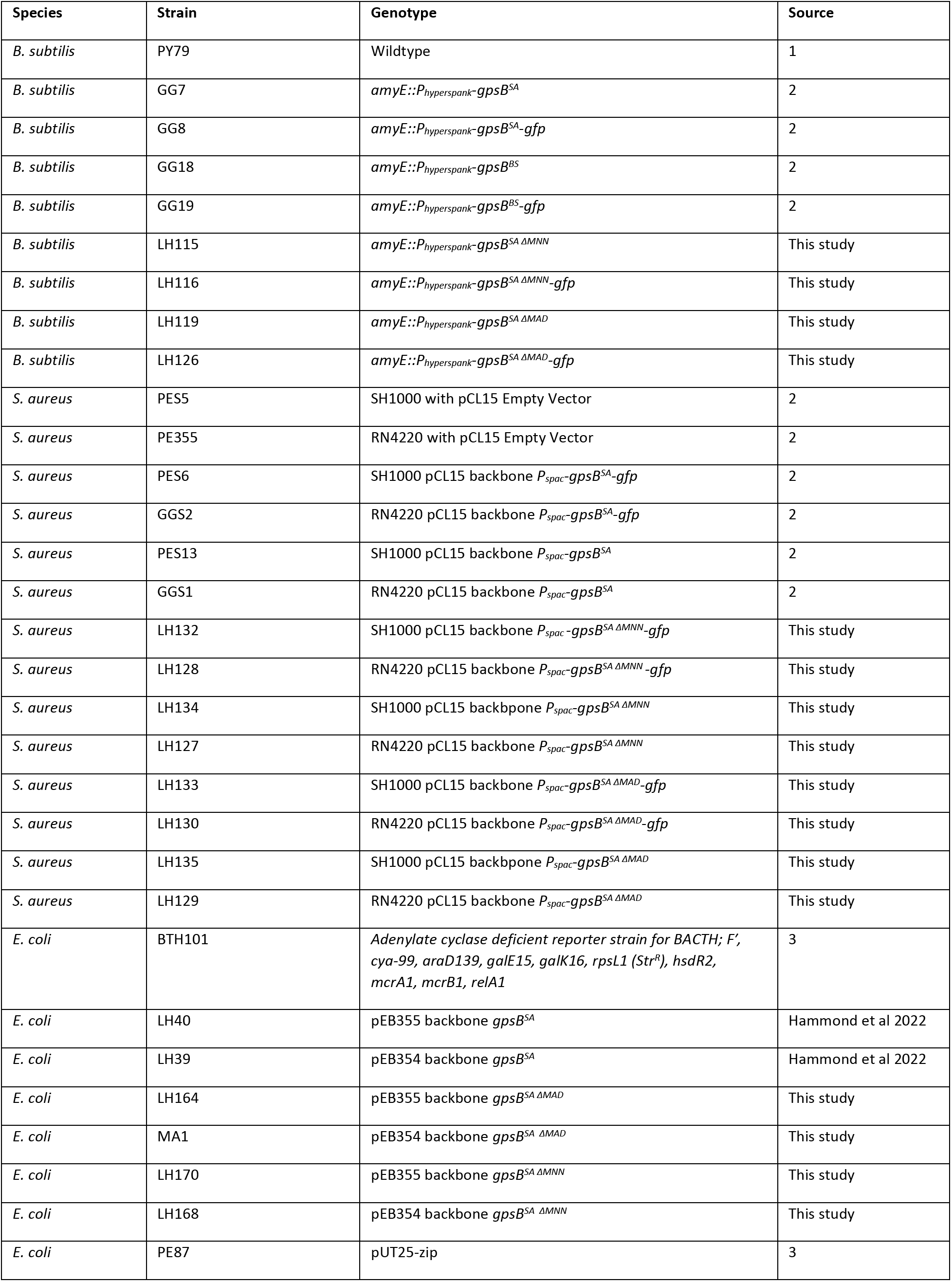
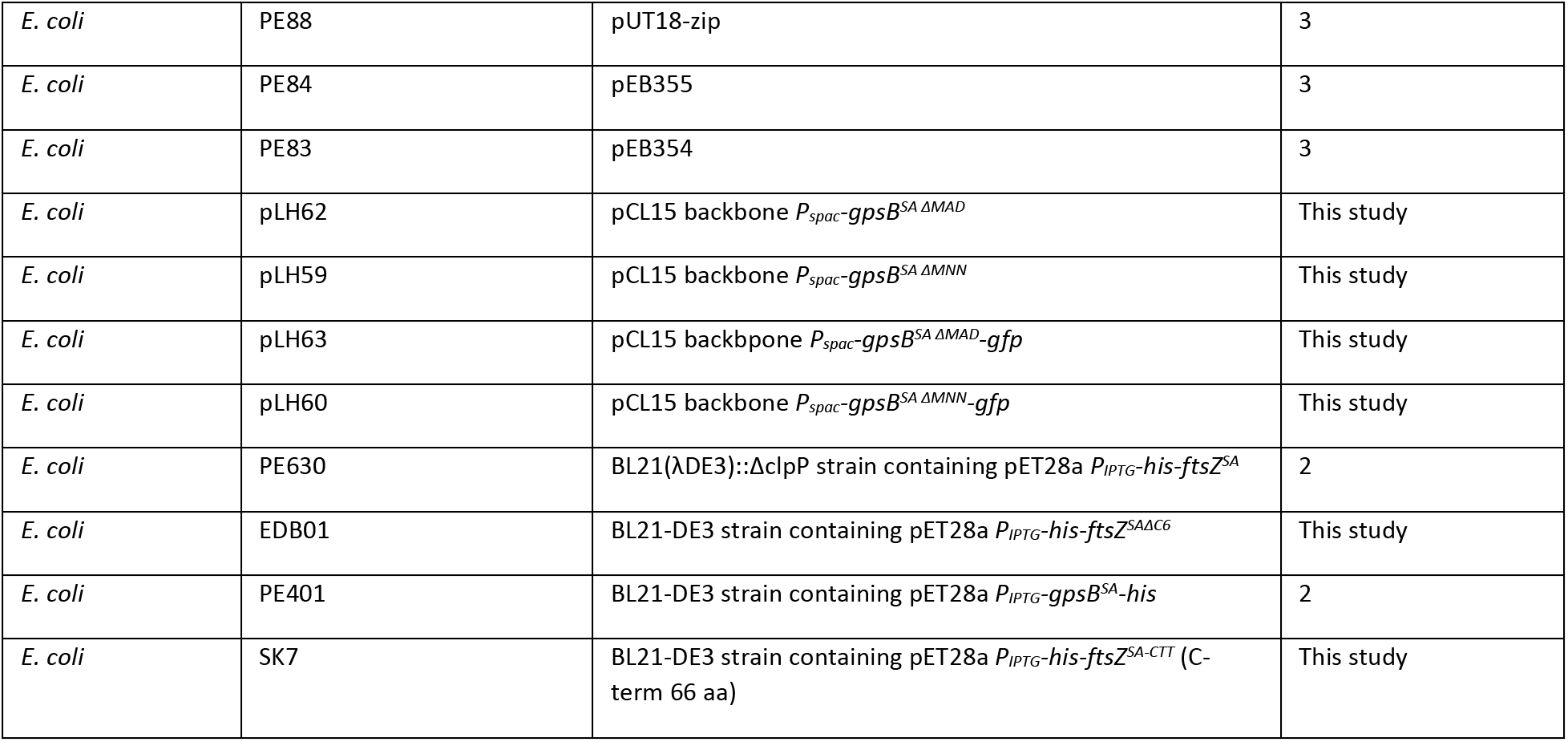
The genotypes of strains used in the cell-based studies.

**Table S4.**
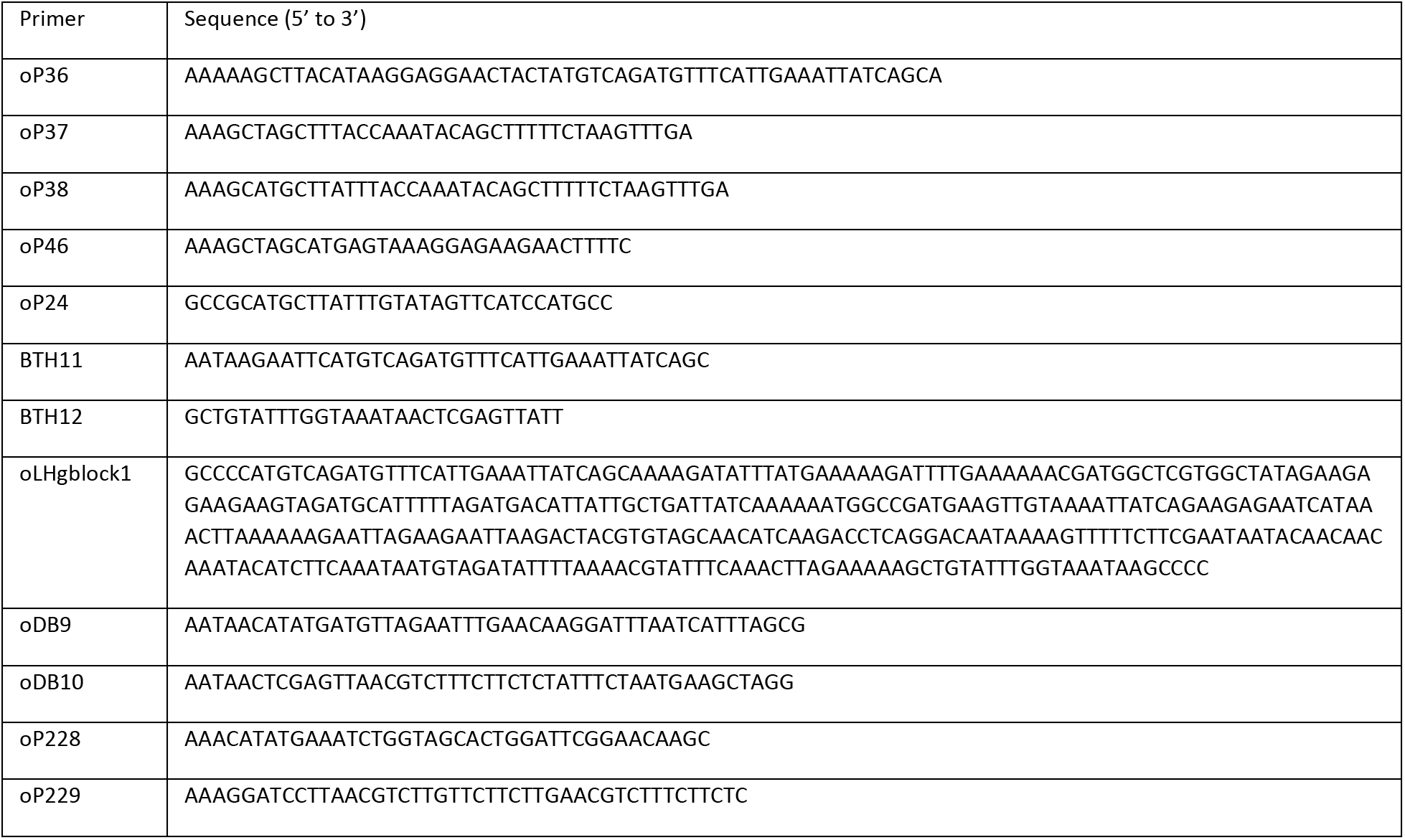
The oligonucleotide sequences used in the cell-based studies.

## Notes

### Competing Interest Statement

The authors have declared no competing interest.

